# Mark-release-recapture of male *Aedes aegypti* (Diptera: Culicidae): use of rhodamine B to estimate movement, mating and population parameters in preparation for an incompatible male program

**DOI:** 10.1101/2020.11.02.365924

**Authors:** Brendan J. Trewin, Daniel Pagendam, Brian J. Johnson, Chris Paton, Nigel Snoad, Scott A. Ritchie, Kyran M. Staunton, Bradley J. White, Sara Mitchell, Nigel W. Beebe

## Abstract

Rapid advances in biological and digital technologies are revolutionizing the population control of invasive disease vectors such as *Aedes aegypti*. Methods such as the sterile and incompatible insect techniques (SIT/IIT) rely on modified males to seek out and successfully mate with females, and in doing so outcompete the wild male population for mates. Currently, these interventions infer the success of mating interactions between male and female insects through area-wide population surveillance and observations of mating competitiveness are rare. Furthermore, little is known about male *Ae. aegypti* behaviours and biology in field settings. In preparation for a large, community scale IIT program, we undertook a series of mark-release-recapture experiments using rhodamine B to mark male *Ae. aegypti* sperm and measure mating interactions with females. We also developed the Spatial and Temporally Evolving Isotropic Kernel (STEIK) framework to assist researchers to estimate the movement of individuals through space and time. Results showed that ~40% of daily females captured were unmated, suggesting interventions will need to release males regularly to be effective at suppressing *Ae. aegypti* populations. Males moved rapidly through the landscape, particularly when released during the night. Although males moved further than what is typically observed in females of the species, survival was considerably lower. These unique insights will lead to a greater understanding of mating interactions in wild insect populations and lay the foundation for robust suppression strategies in the future.

**Author Summary:** Modern scientific techniques for controlling populations of the dengue vector, *Aedes aegypti*, utilize the mating biology of adult male mosquitoes to achieve suppression through a sterilization process. As the study of *Ae. aegypti* control has typically focused on adult female mosquitoes, knowledge on male movement, survival and mating interactions in the field is lacking. Here we undertook several mark-release-recapture experiments on adult male *Ae. aegypti* in Innisfail, Australia, and measured important biological parameters. For the first time in large field experiments, we employed rhodamine B as a marker that when fed to adult males, identified both marked males and the wild females they mated with. We observed males moving further through the landscape, but surviving for a shorter period, than previous measurements undertaken on females in a field setting. A high proportion (~40%) of unmated females suggests individuals are constantly available for mating. As such, sterile male strategies may need to release at regular intervals to achieve effective population suppression. The unique insights provided by this study will assist in designing future sterile male field interventions.

## Introduction

Rapid human population growth and urbanization, combined with widespread resistance to insecticides, have led to a dramatic increase in the incidence of vector-borne diseases such as dengue, chikungunya and Zika (1, 2). In the battle to contain wide-spread epidemics of vector-borne disease, mosquito control has turned to species specific technologies to suppress mosquitoes and the pathogens they transmit, at the landscape scale. Rapid advances in molecular biology, genetics and digital support systems have enabled area-wide ‘rear-and-release’ strategies such as the use of *Wolbachia* induced Cytoplasmic Incompatibility (or incompatible insect technique IIT)(3), the sterile insect technique (SIT)(4) and the *Wolbachia* population replacement method (5). Together rear-and-release strategies are revolutionizing the suppression of mosquito-borne disease as they give rise to the ‘fourth great era of vector control’ (6).

For many decades mark-release-recapture (MRR) studies have been used to understand mosquito movement and population parameters (7). Releasing marked individuals into a population allows for the inference of ecological factors from both released insects and the wild population. Such studies provide estimates of mosquito movement, survival and population size via the Lincoln-Peterson Index (LPI) or its variations (8), all of which have been key to the management of disease spread in the past (9). Traditional mosquito MRR studies have typically focused on adult female movement and ecology because it is this population that drives pathogen transmission (10–12). In contrast, the movement and mating behaviour of male mosquitoes is rarely a major component of MRRs, particularly in *Aedes aegypti* (Linnaeus), one of the world’s most highly studied mosquito species (9).

Understanding male *Ae. aegypti* biological parameters such as survival, dispersal and mating competitiveness, have become increasingly important as the SIT/IIT methods rely on the success of mating interactions. The small number of studies examining male *Ae. aegypti* movement have generally reported variations these parameters. Early movement studies, some utilizing human landing catches, suggest that the majority of male *Ae. aegypti* disperse within 50 m of release sites after a week (12–16). Studies utilizing modern trapping methods have reported similar mean distances travelled which are generally less than female movement measured in the study (12). Interestingly Tsuda et al. (17) measured male mean distance travelled (MDT) to be greater in males than females, while Trewin et al. (18) suggest males move rapidly through the environment, although none were observed to cross movement barriers such as roads. The average life expectancy (ALE) of male *Ae. aegypti* has been estimated to be between 1 and 3 days for both wildtype (13, 17, 19) and transgenic males (20, 21). The final parameter, mating competitiveness between modified and wild strains, is generally inferred from oviposition results in cage or semi-field cage trials (22–24) and rarely in a field setting (25). All three of these biological parameters are essential to the performance of mass-reared male *Ae. aegypti*, affecting the ability of males to efficiently seek out and successfully mate with wild-type females.

The primary challenge for SIT/IIT strategies is the determination of adequate population overflooding across large areas. To do so, one must have a thorough understanding of target population size, demographics and movement within the landscape. Empirically informed models that simulate movement over extended landscapes are cost-effective methods of predicting release efficiency. Standard measures of population dynamics can often be obtained easily enough through traditional surveillance methods, but determination of movement patterns beyond that achieved in limited MRR studies is difficult. Traditionally, movement is classified as summary measures of flight such as a mean or range of the distance travelled and assume movement is a discrete linear distance to traps (18, 26, 27). More recently, mosquito movement studies have incorporated dispersal kernel theory, where distributions of movement can be estimated over the entire flight range (21) and integrate a temporal component such as average life expectancy (18). Accurately parameterizing models for forecasting dispersal is a challenge, primarily due to the lack of accurate data and the expense of collecting these data from field environments. Furthermore, the development of precise models and simulations can only be achieved if field data from multiple ecological and environmental contexts is available to validate results. However, the variety and quality of data obtained is most often dictated by the chosen mark or tracking method.

A variety of marking methods have been employed to infer mosquito movement and behaviour including paints, dyes, trace elements, fluorescent dusts and radioactive and stable isotopes (7). Marking methods are often limited in their effectiveness due to time inefficiencies in application, ability to detect markers, high expense, requirements for technical expertise and physical restrictions imposed by the mark on individual behaviours (7, 28). The fluorescent dye rhodamine B is a recent innovation in the use of fluorescent markers to stain male spermatophores in insects and has provided a rapid and cheap way to understand mating interactions (29–33). Rhodamine B provides field ecologists with a method to measure both movement and mating interactions through the staining of male sperm, seminal fluids and body tissues. Producing a distinct bright red colour fluorescenece when excitated under 540 nm (maximum excitation) and 568 nm (maximum emission) light wavelengths, the dye can be observed in ~ 95% of female *Ae. aegypti* spermathecae after four days (29). The method allow investigators to mark both male and female mosquitoes, determine key performance indicators and rapidly infer the efficacy of an intervention by measuring behavioural and ecological factors such as mating success.

Thus, in preparation for a large, community-scale rear-and-release IIT program (Debug Innisfail, Australia, 2017-2018) we aimed to quantify the movement and mating behaviour of male *Ae. aegypti* through urban landscapes in north Queensland. To do this we undertook a number of rhodamine B-based MRR experiments, utilizing wild-type male *Ae. aegypti* to examine key biological parameters across a number of spatial, temporal and climatic scenarios.

## Methods

### Study Sites

Six mark-release-recapture experiments were run during two seasonal periods in North Queensland, Australia. Mark-release-recapture experiments 1-3 (season 1) occurred before the wet season, between the 18^th^ November 2016 and 13^th^ of December, while MRR experiments 4-6 (season 2) occurred during the wet season between the 7^th^ and 27^th^ of February 2017. The study site in South Innisfail (17.5435 °S, 146.0529 °E) was situated to the south east of Innisfail, a rural town on the main highway 88 km south of Cairns, in a residential area, 0.18 km^2^ in size. The site contained 95 residential premises bounded by the Johnson River to the West and by grass sports fields and forest to the east. The site also contained a primary school to the north and a number of small commercial buildings (Figure 1). The Innisfail region is one of the wettest in Australia, averaging 3,547 mm of rainfall annually with tropical cyclones occurring throughout the Summer and Autumn (34). The urban landscape of Innisfail is unusual for northern Australia, with dwellings in the town a mix of Queenslander (constructed of wood with tin rooves and typically raised off the ground by 1.5-2 m) single floor fibre board, modern brick single floor, and ‘art deco’ style single floor constructions. House block size were approximately 800 m^2^ with simple fencing or hedge-like greenery on boundaries, with open space underneath raised buildings utilized for storage, laundry and recreation areas. Roads averaged 25 m wide (fence to fence).

**Fig 1.**
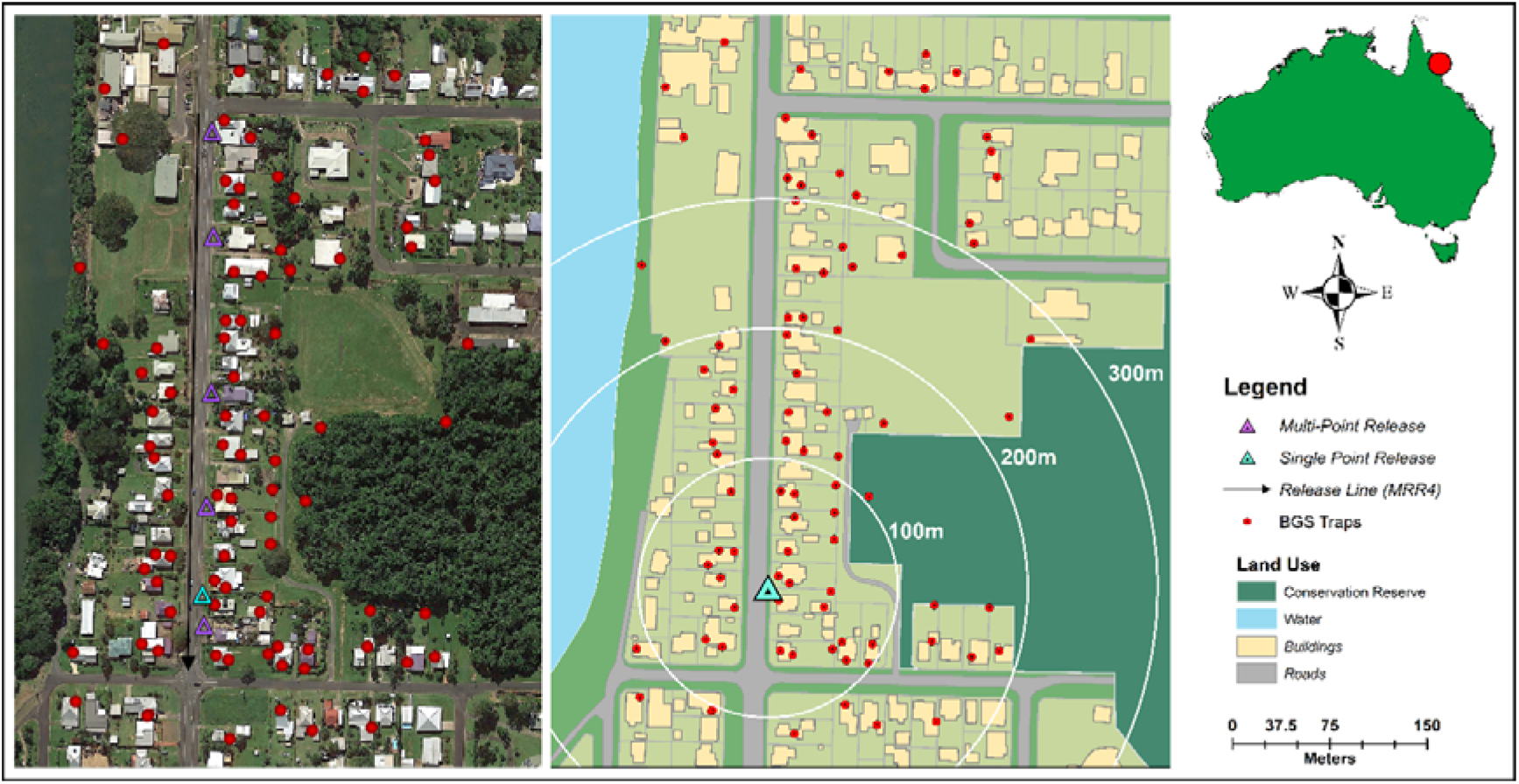
Location of study site in South Innisfail, Australia. Maps indicate landscape characteristics which include natural topography (left) and land use (right). Rhodamine B marked *Aedes aegypti* were released at single point (blue triangle) and multipoint locations (purple triangles) and recaptured using Biogents Sentinel traps (red circles). Base layer of region sourced from Google Earth 7.3.3.7786. (July 21, 2020). South Innisfail, Australia. 17.5435 °S, 146.0529 °E, SIO, NOAA, U.S. Navy, NGA, GEBCO. TerraMetrics 2012, DigitalGlobe 2020. http://www.earth.google.com [November, 2020](35).

### Rearing and Release

For colony establishment, wild type *Ae. aegypti* were sourced from multiple ovitrap collections and locations in Innisfail during 2016. Larvae from field collected ovistrips were hatched, sorted to species, and reared to adults. All mosquito colonies were maintained using standard laboratory rearing protocols at 28 °C ± 1 °C, a 70 % (± 10 %) relative humidity with a 12:12 hour light cycle and twilight period. The adult colony was supplemented with regular ovitrap collections from Innisfail leading up to MRR experiments and maintained at a population size of 300 individuals.

Wild-type *Ae. aegypti* eggs were hatched into a 0.2 g/L yeast in water and allowed to feed for 24 hours. Five-hundred first and second instar larvae were pipetted into a three-litre bucket to an approximate density of one larvae per 6 ml of water. Larvae in each bucket were reared on ground Tetramin Tropical Fish Flakes (Tetra, Germany) and provided with 0.45 g on day 2, 0.8 g on day 5 and again on day 6 if required. Ten minutes after food settles, bucket water was stirred in a ‘side to side’ motion to distribute ground fish flakes evenly throughout the rearing buckets. Males were separated with a one ml bulb pipette based on pupal size estimated by eye and 20 individuals removed via pipetting into meshed, 300 ml Styrofoam rearing cups. After emergence, cups were visually inspected for the presence of females and if detected these were removed through aspiration. Adult males were fed a 0.4 % rhodamine B (weight to volume) solution. The solution consisted of 160 mg rhodamine B dissolved in 40 ml of a 25 % honey solution following the methods of Johnson, Mitchell (29). Males were maintained on the solution for four days to ensure adequate body and seminal fluid marking (29). Males were transported to study sites the day before release and released at five days of age in all experiments.

Approximately 1,250 males were released during each of the six MRR experiments. Releases occurred at 6am for day releases for MRR 1-5 and 7 pm for the night release (MRR 6). Release location varied depending on experimental design, with single point releases occurring at the southern end of the study site (MRR 1, 2 & 6), multi-point releases occurred at five points along the release block (MRR 3 & 5) and a single linear release (MRR 4) from a prototype of the mechanical device used in Crawford et al. (36). This involved a consistent linear release of males along the eastern side of Mourilyan road from north to south (Figure 1).

### Trapping Arrays and Recaptures

The study site contained 83 Biogents Sentinel traps without lures (BGS; Biogents GmbH, Regensburg, Germany) set with the goal of distinguishing landscape characteristics that affected the capability of males to move through blocks and across movement barriers such as roads (Figure 1). To do this, one trap was placed close to each chosen dwelling, one in the backyard and, where possible, the forested area adjacent to the residential area. Additional traps were placed at dwellings across the road from release sites to monitor movement across a known dispersal barrier. For releases one and two, collections began one day post-release to allow for unbiased mixing with wild population. For MRRs 3-6, traps were turned on two hours post-release.

All traps were serviced daily throughout each MRR upon activation. Captured adult *Ae. aegypti* were stored at ~4 °C for transfer to the laboratory for identification and both males and females processed for the presence of rhodamine B following the methods of Johnson et al. (29). Females were considered mated by a released marked male if rhodamine B was observed in the bursa, spermathecae or both. Females were considered to have mated with a wild, unmarked male if sperm, visualised by DAPI staining, was present in the bursa, spermathecae or both in the absence of rhodamine B.

### Determination of Biological Parameters, Statistical Analysis and Dispersal Kernel Framework

For all experiments the PDS was estimated by regressing log_10_ (x+1) the number of recaptured males against the day since release where the antilog_10_ of the regression slope is the PDS (37). The average life expectancy is then calculated from the PDS as 1/-log_e_PDS (38). Population estimates, where applicable, were calculated via the LPI:

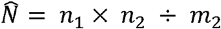

Where 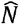 is the size of the population, *n*_1_ is the number of marked animals released into the population, *n*_2_ the total number of individuals captured and *m*_2_ the total number of marked individuals recaptured. All five assumptions of the LPI were met (8) which include: 1) the mark should not affect insects, 2) marked insects are allowed to become completely mixed with the local population, 3) sampling is random with respect to marked insects, 4) samples are measured at discrete time intervals in relation to total time, and 5) the population is not unduly influenced by immigration or emigration during the period of study. Traditional methods of MDT were calculated using the methods of Lillie et al (27) and Morris et al (26) where:

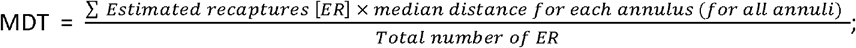

and applying a correction factor (CF) to accommodate unequal trapping densities

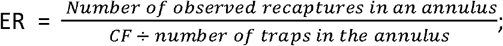

where

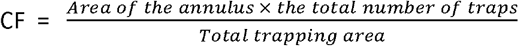

Flight range (FR) of male movement are estimated from the linear regression of the cummulative ERs for each annulus on the log_10_ as the value of *Y* at 50% and 90% (13). We introduce the concept of mean insemination distance (MID) by modifying the above methods of Lillie et al (27) and Morris et al (26) by estimating the mean distance over which rhodamine B inseminated females were captured during each experiment.

We then compared traditional MDT estimates with the spatially and temporally evolving isotropic kernel (STEIK) framework developed by Trewin et al. (18). The STEIK framework uses an isotropic Gaussian diffusion model with kernels defined as a temporally-evolving probability density function (PDF) over two-dimensional space (18). Here the probability of a mosquito being trapped per unit area is a function of the distance from the release location and the time since release (18). For the multi-point releases in this study we divided the total number of mosquitoes released evenly between release points. For STEIK estimates of 50% and 90% FR, quartiles of simulated kernel distributions were calculated from parameter estimates relevant to average lifetime and the standard deviation of the isotropic kernel for each experiment. To facilitate the use of our STEIK framework by experimentalists, we have stored male recapture data and R code at https://github.com/dpagendam/MRRk (39). Raw mosquito and trap data is available at https://doi.org/10.25919/5f9f642323d86.

To examine differences in the daily proportion of rhodamine B inseminated females with the total number of mated females between seasons, we used a mixed effects, logistic regression model with a binomial distribution and logit link function. Fixed effects included season and release type (multipoint vs point release) with a random effect of experiment number. The same model framework was used to examine differences in the total daily proportion of mated females (both rhodamine B and wild mated) between seasons and the daily proportion of wild-type mated females with total females captured. The Akaike information criterion (AIC) was used to selecte the most parsimonious model. Odds ratios (OR) were calculated for coefficients exhibiting significant differences in proportions. The R package ‘glmmTMB’ was used for all mixed effects models and the packages ‘DHARMa’ was used for model diagnostics and ‘ggplot2’ for visualisations. To look for collinearity in predictors, correlations were examined using the R package ‘corrplot.’ To compare whether rhodamine B and wild males and wild female *Ae. aegypti* were more likely to be captured by BGS traps at certain locations (house, backyard or forest), we used contingency table analysis with odds ratios calculated via the R package ‘epitools’. All analyses were performed using R version 3.5.3 (40). All landscape maps were digitized by outlining landscape features (houses, roads, blocks, river) in Google Earth (35) and modified in ArcGIS Desktop. Two-dimensional kernels were output as images from an R density function where the mean was equal to zero and the spread equal to the time dependent standard deviation of the kernel. Kernels were then overlaid on maps to scale in standard image editing software.

### Ethics and Community Engagement

Human ethics was sought through the CSIRO Social and Interdisciplinary Science Human Research and Ethics Committee (CSSHREC) and approved under project 026/16 named “Sterile insect technology development for *Aedes aegypti*”. As part of this approval all residents in release areas provided voluntary consent for scientists to operate within their property, and were provided with an information sheet detailing how, why, where and when the research was to be performed and funding bodies. All residents were informed about the risks and benefits, including the potential for an increase in mosquito numbers during male releases. To enhance communication, brochures were distributed to homeowners, articles were posted in local newspapers, a website was setup for enquiries and residents were engaged through a project advisory group containing members of the local community.

## Results

### Population Statistics

Environmental conditions varied considerably both during and before each season. The most notable difference being total rainfall 2 weeks before each season, with combined totals of 65.8 mm and 762 mm falling before MRRs in season 1 and 2, respectively (Tables S1)(34). Mean daily minimum and maximum temperatures during the study periods varied between 15.1 °C and 31.9 °C, and the mean relative humidity at 09:00 and 15:00 hours varied from 85.7 % (SD ± 6.8) and 53.8 % (SD ± 6.8), respectively (Tables S1). Wind direction was predominantly from the South East (Figures S2). Total rainfall for the two weeks leading up-to-and during releases varied from 0 mm in November to 344 mm in February (Tables S1)(34). Visual assessment of wind direction and speed suggest it played little part in the direction of daily male *Ae. aegypti* movements (Figures S3).

A total of 313 (4.1 %) male *Ae. aegypti* were recaptured from a total of 7,713 released into the South Innisfail landscape. Recapture rates for rhodamine B marked *Ae. aegypti* males varied during individual experiments (0.7 % to 10.4 % recaptured), while the mean recaptures increased on day two (mean = 24.5, SD ± 22.0), before decreasing (Table 1, Figure 2). Recapture success was highly variable primarily due to time of year, with the early summer and later summer averaging 1.0 % (SE ± 0.2) and 7.3 % (SE ± 4.3) recaptured, respectively (Table 1). Maximum time to recapture (the period between release and last date of a marked individual captured) varied from three to seven days across all experiments (Table 2). There were significantly more wild male and female *Ae. aegypti* caught daily in season two that season one (Figure 3, *F*_(2,41)_ = 18.01, P = <0.001).

**Fig 2.**
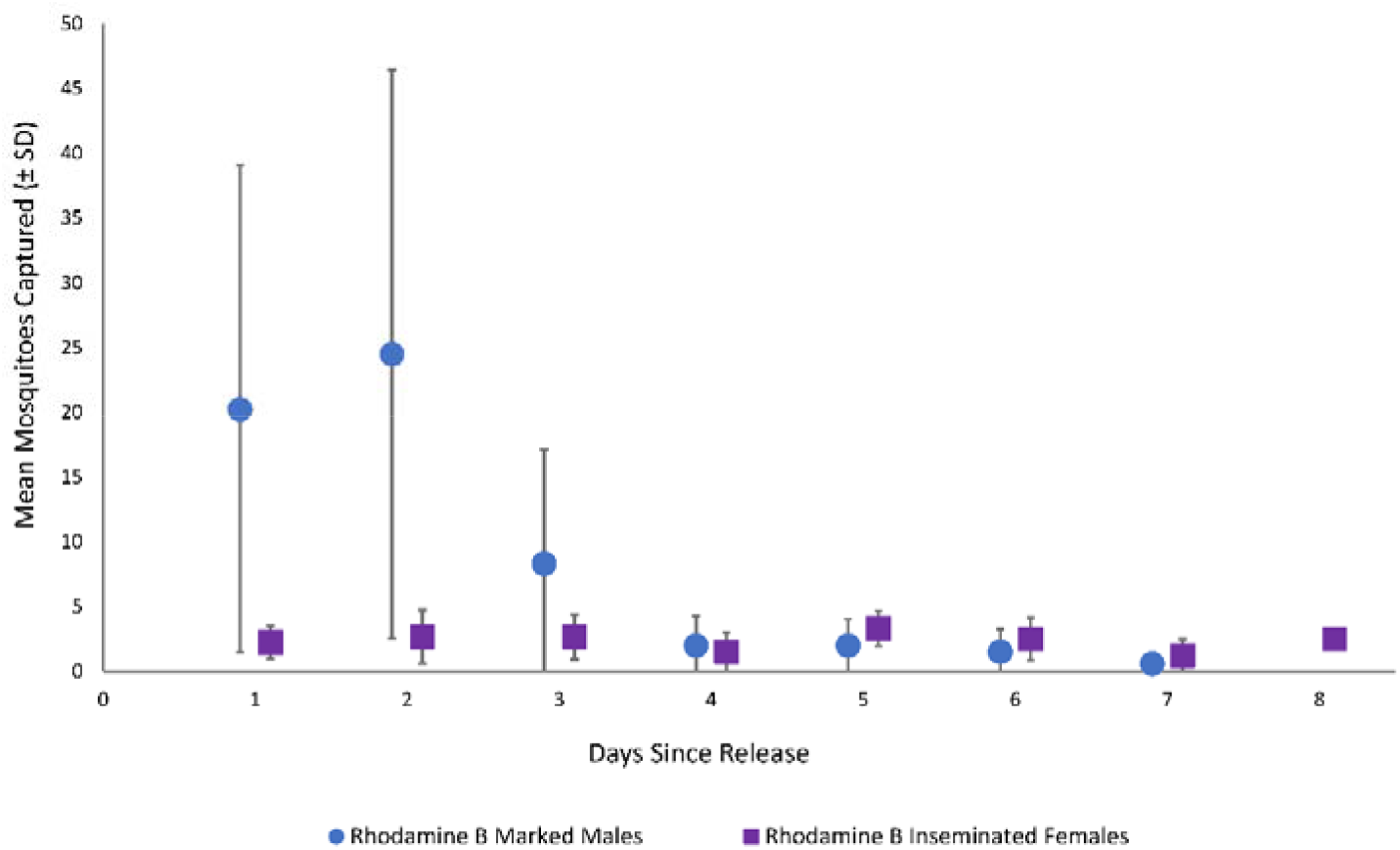
Daily mean recapture rate (SD) of rhodamine B marked male (blue circles) and inseminated female *Aedes aegypti* (purple squares). Data aggregated across all mark-release-recapture experiments in South Innisfail, Australia.

**Fig 3.**
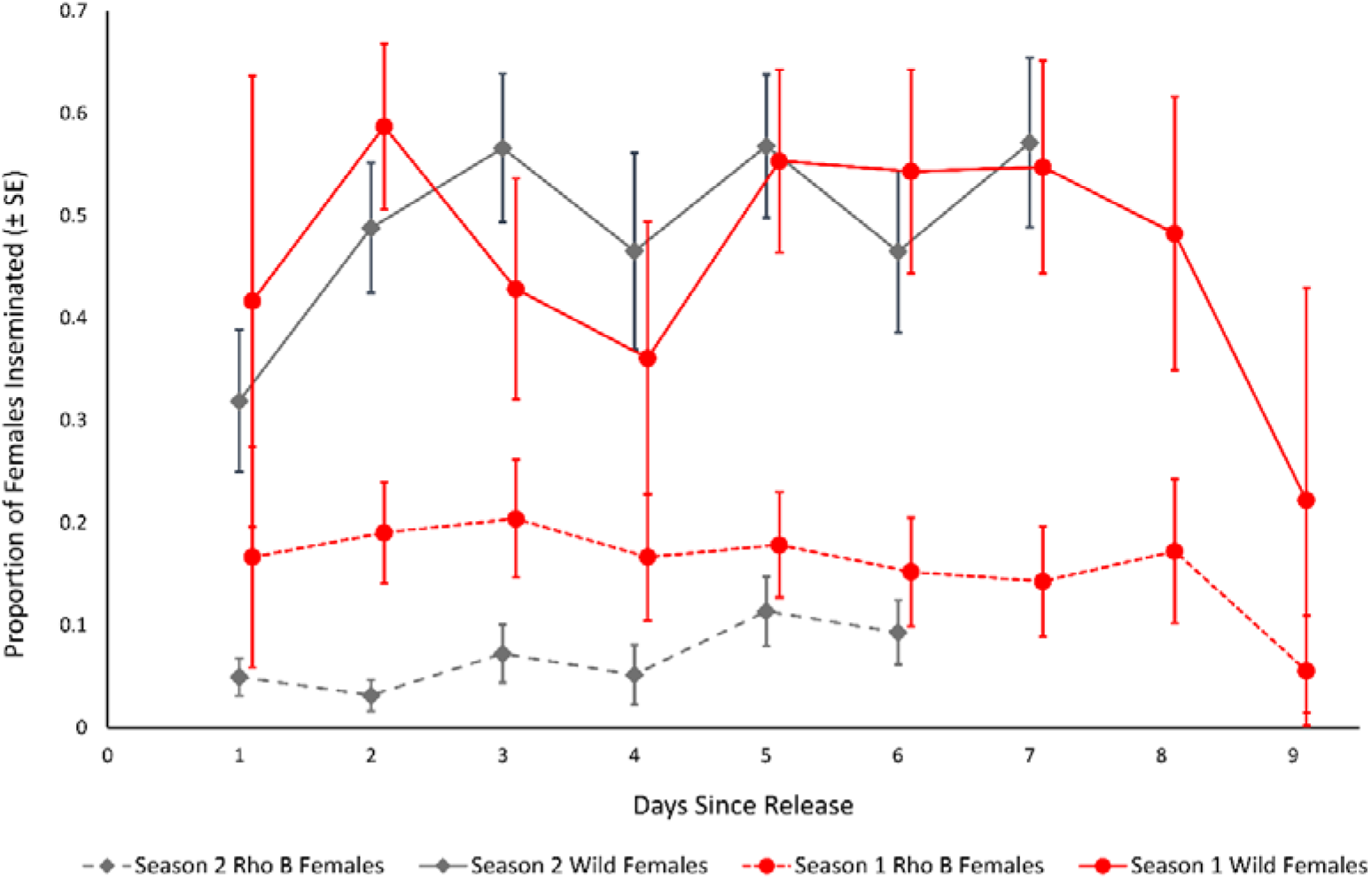
Daily proportion (SE) of captured *Aedes aegypti* females inseminated by wild or rhodamine B marked males. Differences between time of year are indicated by early summer (season 1, red circles) and late summer (season 2, grey diamonds).

**Table 1.** Mating and recapture results from individual rhodamine B marked *Aedes aegypti* experiments in South

|  |  | Rho B |  | Total | Rho B | Females |  |  | Innisfail, Australia. Wild female s were examined for rhoda |
| --- | --- | --- | --- | --- | --- | --- | --- | --- | --- |
|  | Rho B | Males | Wild | Males | Mated | Mated by |  | Total |  |
|  | Males | Recaptured | Males | Captured | Females | Wild Males | Unmated | Females |  |
| Release | Released | (%) | Captured | (% Rho B) | (%) | (%) | Females | (% Mated) |  |
| 1 | 1228 | 17 (1.4) | 72 | 89 (19.1) | 22 (18.9) | 61 (52.6) | 33 | 116 (71.6) |  |
| 2 | 1485 | 11 (0.7) | 51 | 62 (17.7) | 28 (17.4) | 82 (50.9) | 51 | 161 (68.3) |  |
| 3 | 1240 | 12 (1.0) | 28 | 40 (30.0) | 9 (12.2) | 30 (40.5) | 35 | 74 (52.7) |  |
| 4 | 1250 | 42 (3.4) | 130 | 172 (24.4) | 13 (4.4) | 127 (43.2) | 154 | 294 (47.6) |  |
| 5 | 1250 | 130 (10.4) | 57 | 187 (69.5) | 13 (5.4) | 123 (51.5) | 103 | 239 (56.9) |  |
| 6 | 1250 | 102 (8.1) | 42 | 144 (70.8) | 12 (10.6) | 57 (50.4) | 44 | 113 (61.1) |  |

mine B insemination by fluorescent microscopy.

**Table 2.**
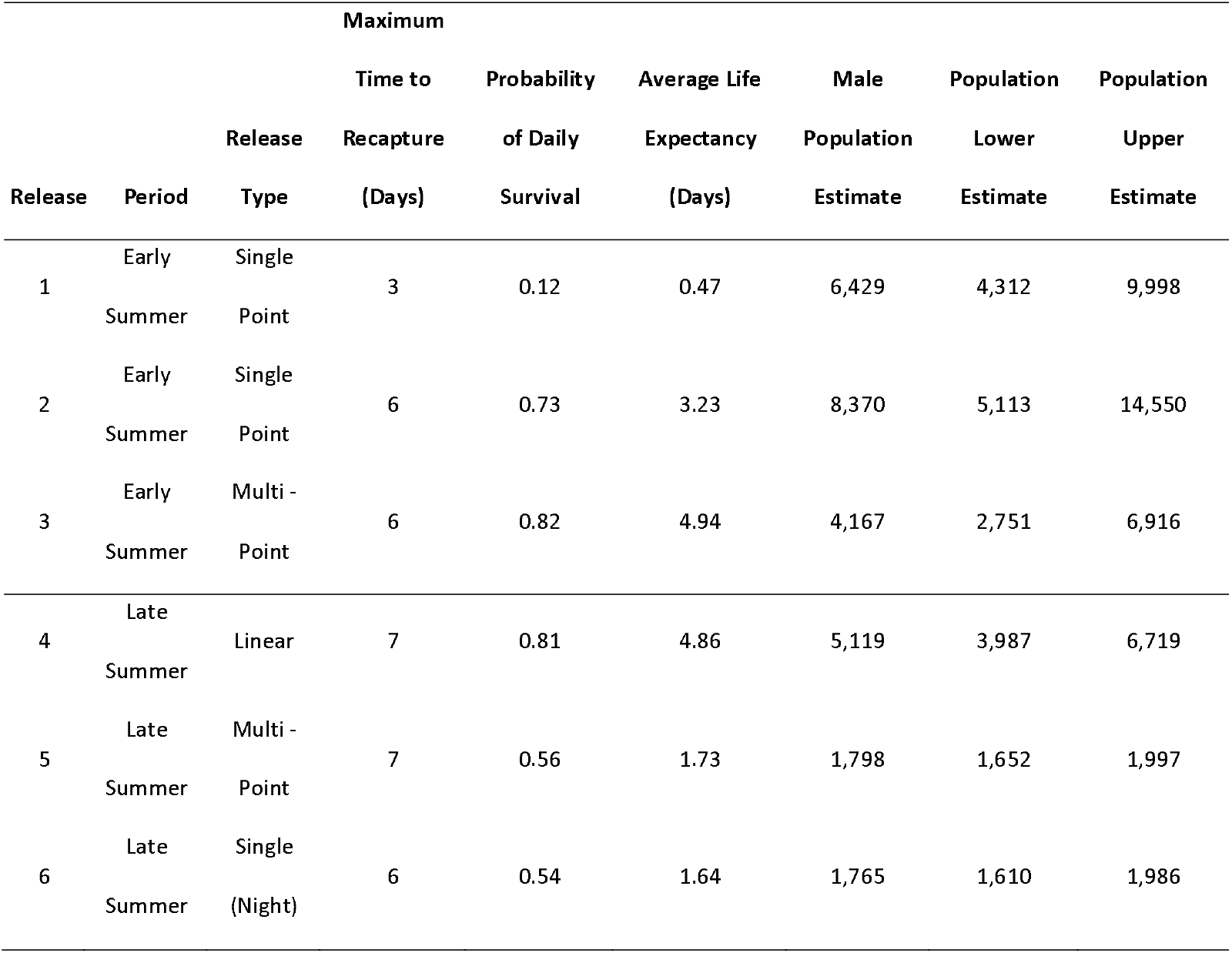
Probability of daily survival, average life expectancy and population estimates from rhodamine B marked *Aedes aegypti* during six mark-release-recapture experiments in South Innisfail, Australia. Wild male population sizes were estimated via the Lincoln Peterson Index (8) and probability of daily survival and average life expectancy via the methods of Gillies (37) and Niebylski and Craig Jr (38).

For estimates of survival and ALE for each experiment, the total number of recaptured rhodamine B marked male *Ae. aegypti* were plotted against days since release (Table 2). When combining data across all MRR experiments we estimated the ALE as 1.69 days (daily survival probability = 0.55, R^2^ = 0.9). The maximum ALE observed in an individual experiment was 4.9 days (daily survival probability = 0.82, R^2^ = 0.99) during MRR 3, and a minimum of 0.47 days (daily survival probability = 0.12, R^2^ = 1) during MRR1 (Table 2). Estimated male population sizes ranged from 1,765 (95 % CI Low = 1,610, High = 1,986) to 8,370 (Table 2; 95 % CI = Low 5,113, High 14,550) with a mean of 4,608 (SD ± 2,604).

### Mating Interactions

The daily capture rate of rhodamine B inseminated females tended to be low but consistent across all experiments (Figures 2, 3 and 4). Likewise, the proportion of all inseminated females (wild and rhodamine B) remained relatively constant across experiments, while total rhodamine B inseminations tended to vary relative to season and total females captured and between 25-52% of females captured per experiment remaining unmated (Figures 3 and 4). The mixed effects logistic regression model revealed the daily proportion of females inseminated by rhodamine B marked males was significantly higher in season one than season two (Z = −2.81, df =37, *P* < 0.005) and in single point than multipoint releases (Figure 5; Z = −2.39, df = 37, P = 0.017), respectively. This equated to a ~54 % and ~48 % decrease in the daily odds of a female being inseminated by rhodamine B marked males during season two (Table 3; OR = 0.46, 95 % CI Low = 0.27, High = 0.79) and during linear releases (OR = 0.52, 95 % CI Low = 0.30, High = 0.89), respectively. However, when the same statistical model was used to examine the proportion of daily total mated females to total female mosquitoes captured, there was a significantly higher proportion in single point releases (Z = −2.876, df = 38, P < 0.004) but not season (Z = −1.20, df = 38, *P* =0.23). Furthermore, there was no significant relationship between the proportion of wild type male *Ae. aegypti* inseminations to total females captured between seasons (Z = −0.39, df = 38, P = 0.93).

**Fig 4.**
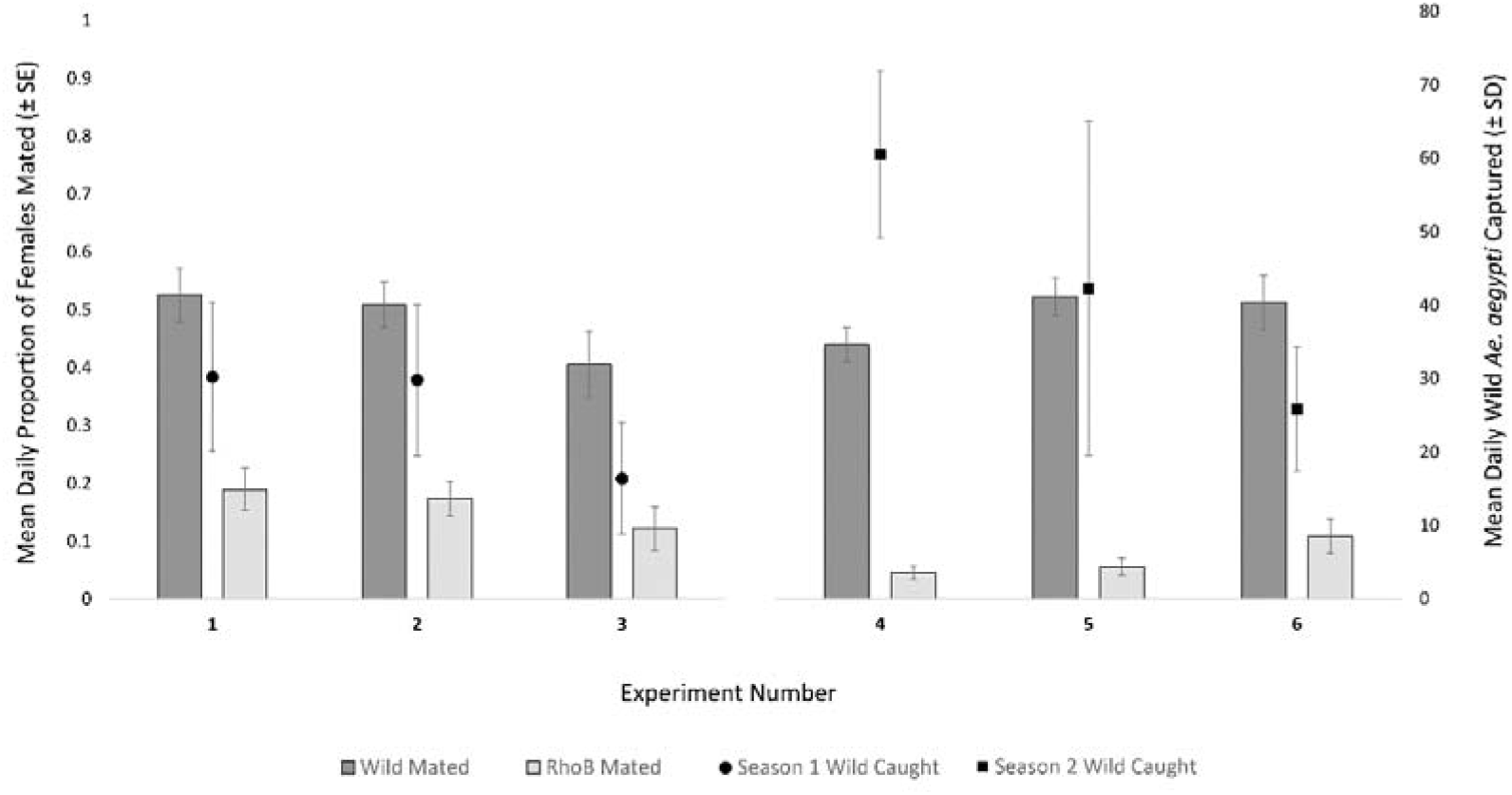
A comparison of mating rate with wild *Ae. aegypti* captured during six mark-release-recapture experiments in Innisfail, Australia. Primary axis indicates the mean daily proportion (± SE) of wild and rhodamine B inseminated female *Ae. aegypti*. Second axis indicates the mean daily total (male and female) number (± SD) of wild male and female *Ae. aegypti* captured during each experiment.

**Table 3.** Results of the mixed effects logistic regression model on female mating rates. Results show the effect of release type (single or multi-point) and season (early and late summer) on the daily proportion of rhodamine B inseminated females captured.

| Fixed Effects | | $\beta$ | $SE \beta$ | z value | $P$ | odds ratio |
| --- | --- | --- | --- | --- | --- | --- |
| Constant |  | -1.49 | 0.15 | -9.82 | <0.001 | 0.23 |
| Release Type |  | -0.66 | 0.28 | -2.39 | 0.017 | 0.52 |
| Season |  | -0.77 | 0.28 | -2.81 | 0.005 | 0.46 |
| Random Effect | | $\sigma^2$ | $SE \beta$ | | | |
| Experiment |  | <0.001 | <0.001 |  |  |  |
| Overall Model |  |  |  |  |  |  |
| Evaluation | Test |  |  |  |  |  |
| | Kolmogorov -Smirnov | | | D = 0.149 | $P = 0.291$ | |
| | Overdispersion | | | Ratio = 0.878 | $P = 0.332$ | |

### Movement Estimates

Males moved rapidly through the environment and were observed up to 422m from the single point of release. Movement was highest during the night release experiment for all metrics calculated (MRR 6; Table 4). The multi-release point isotropic Gaussian kernel framework provided MDT estimates comparable to those of Lillie, Marquardt (27). When MDT is estimated over the entire period of each experiment, the STEIK generally estimated larger movement distances than the traditional method. For example, males were estimated to travel a mean of 295.2 m (FR_50/90_248/480 m) compared with 451 m (FR_50/90_335/873 m) by traditional and STEIK methods, respectively (Table 4). The highest STEIK MDT estimate was 462.8 m (Figure 6; FR_50/90_326/837 m) compared to a traditional estimate of 310.5 m during MRR6 (Table 4; FR_50/90_261/508 m). Male MDT during the multi-point releases in MRR3 and MRR5 were unable to be estimated using the traditional method of Lillie et al. (1981), as it relies upon a discrete point of release. However, the multi-point, STEIK framework estimated an MDT of 110.4 m (FR_50/90_76/205 m) and 184.8 m (FR_50/90_ 132/333 m) for MRR3 and 5, respectively (Figure 5, Table 4). The MDT for MRR4 was unable to be calculated by either traditional or multi-point isotropic kernel methods as this release did not use discrete points as male releases were by a moving vehicle and regarded as linear. Using average life expectancy to estimate a distribution of survival, estimates of MDT were generally lower or equal to those estimated by the isotropic kernel or traditional methods that incorporate the maximum time to recapture (Table 4). The maximum distance over which rhodamine B inseminated females were captured was greater than marked males in MRRs 1 & 2, with marked females caught at the maximum distance of our trapping network. Total and daily female MID were considerably higher than male movement calculations (Table 5).

**Fig 5.**
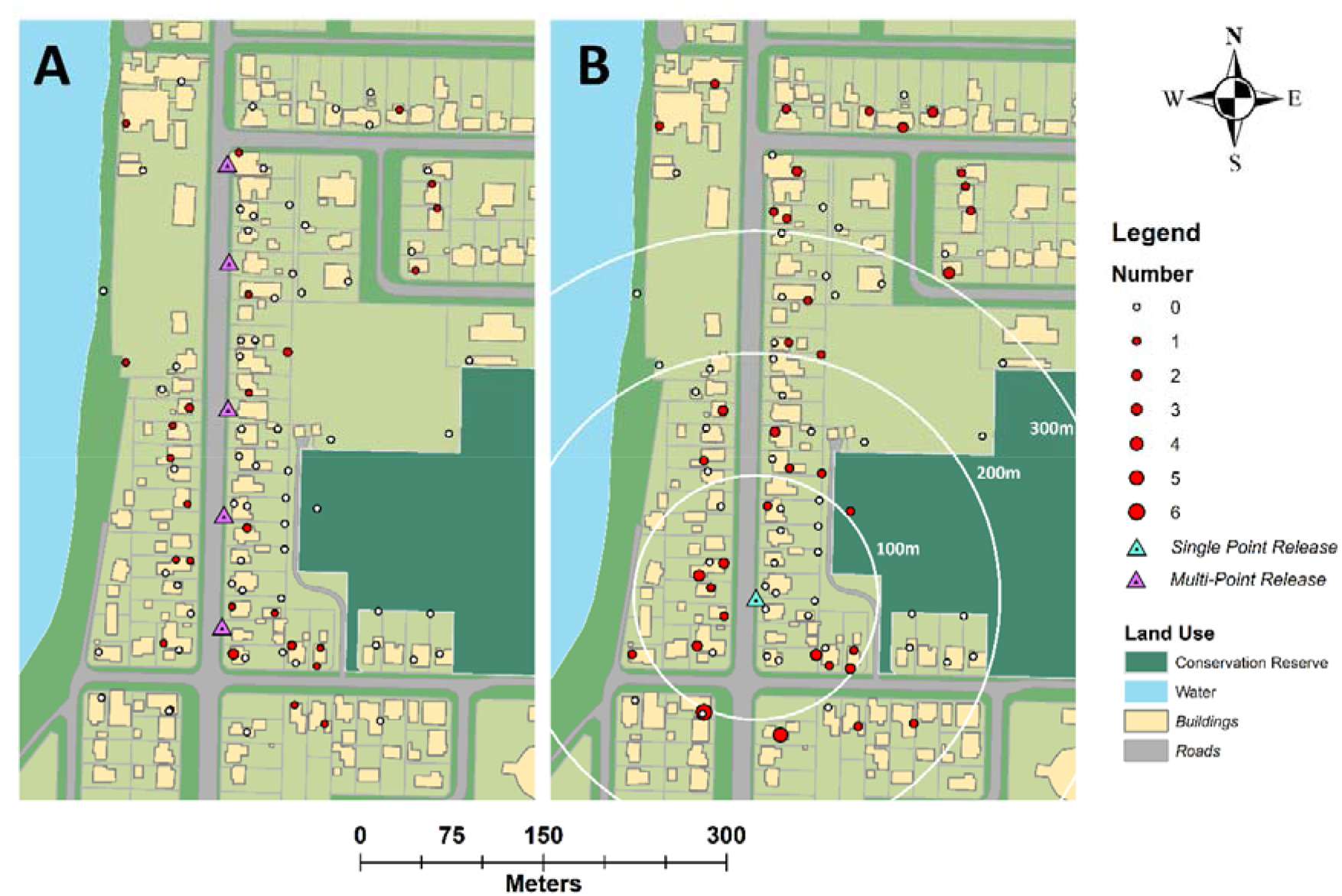
Total rhodamine B inseminated Aedes *aegypti* females captured during multipoint (A) and single point releases (B). Size and colour of circles indicates the total females caught in an individual trap (see key). Release points for each release type are indicated by blue triangle (single point) and purple triangles (multi-point).

**Fig 6.**
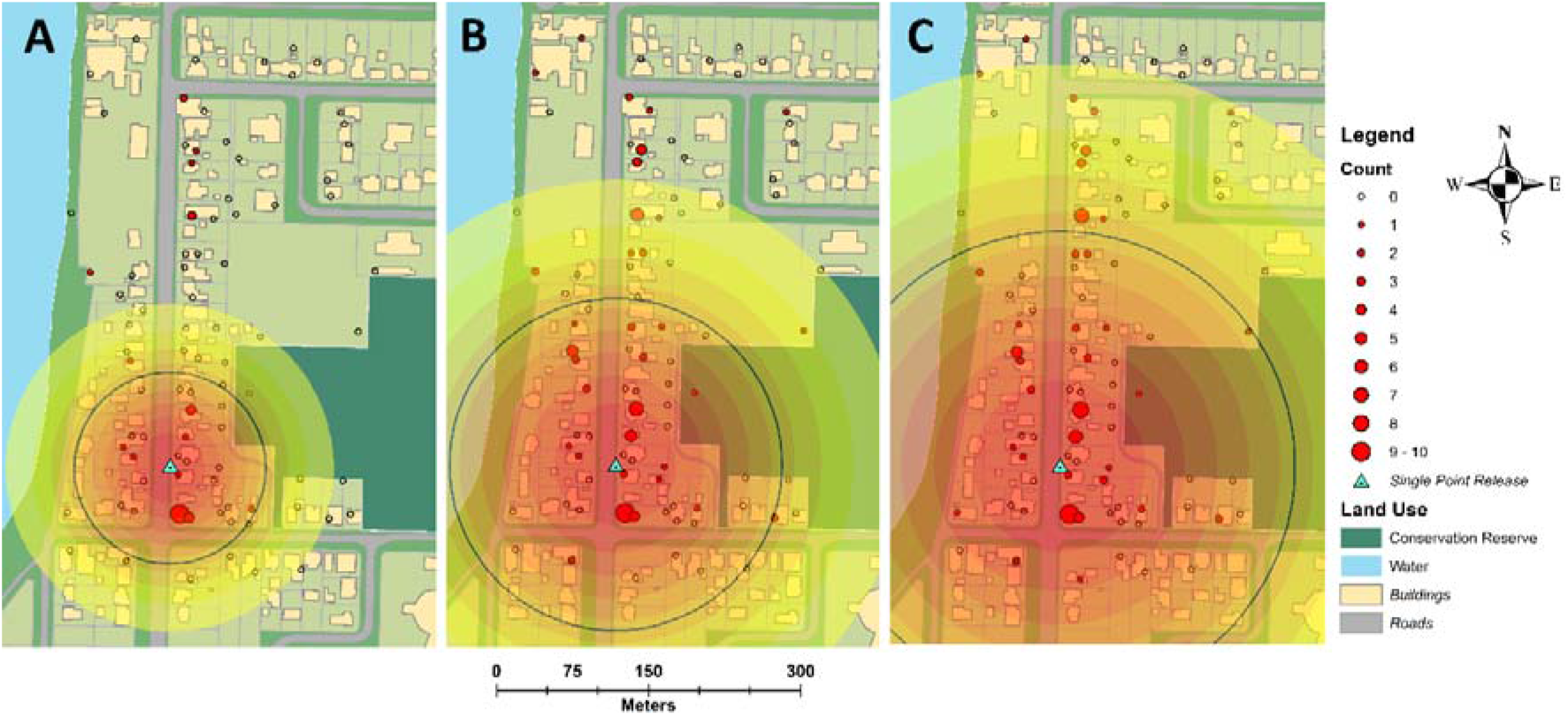
Single point estimates of *Aedes aegypti* movement during experiment six which apply the STEIK framework. Concentric circles from release point are density estimates of marked adult male *Aedes aegypti* one (A), three (B) and six (C) days post release. Black lines represent mean distance travelled over the time period.

**Table 4.** Movement estimates Male *Aedes aegypti* from six mark-release-recapture experiments in South Innisfail, Australia. Spatially and Temporarily Evolving Isotropic Kernel (STEIK) framework compared with the traditional annulusbased methods. The STEIK framework allows for estimates of movement based on average life expectancy and from multiple release points (MRR 3 & 5).

| MRR | Maximum |  |  |  |  |  |  |  |  |
| --- | --- | --- | --- | --- | --- | --- | --- | --- | --- |
|  | Distance | Max | Traditional | Flight Range | STEIK | Flight Range | Average Life | STEIK | Flight Range |
|  | Travelled | Time | MDT | 50/90% | MDT | 50/90% | Expectancy | ALE-MDT | 50/90% |
|  | (m) | (d) | (m)<sup>β</sup> | (m) | (m)* | (m) | (d) | (m)‡ | (m) |
| 1 | 187 | 3 | 126.7 | 118/261 | 95.4 | 68/172 | 0.5 | 51.8 | 43/109 |
| 2 | 282 | 7 | 233.6 | 202/342 | 278.3 | 195/499 | 3.2 | 179.6 | 146/367 |
| 3 | na | 6 | na | na | 110.4 | 76/205 | 4.9 | 95.0 | 75/195 |
| 4 | na | 7 | na | na | na | na | 4.9 | na | na |
| 5 | na | 7 | na | na | 184.8 | 132/333 | 1.7 | 93.5 | 76/187 |
| 6 | 422 | 6 | 310.5 | 261/508 | 462.8 | 326/837 | 1.6 | 246.7 | 199/501 |
| Mean | 297 | 6 | 295.2 | 248/480 | 451.7 | 335/873 | 1.6 | 240.8 | 195/497 |

<sup>β</sup> Calculated using the traditional methods of Morris, Larson (26) and Lillie, Marquardt, and Jones (1981) over the period of study.
\* Method for MDT uses a random lifetime generated from the maximum time to recapture.
¥ Method samples from one million lifetimes with a distribution equivalent to the estimated daily survival rate from the field.

**Table 5.**
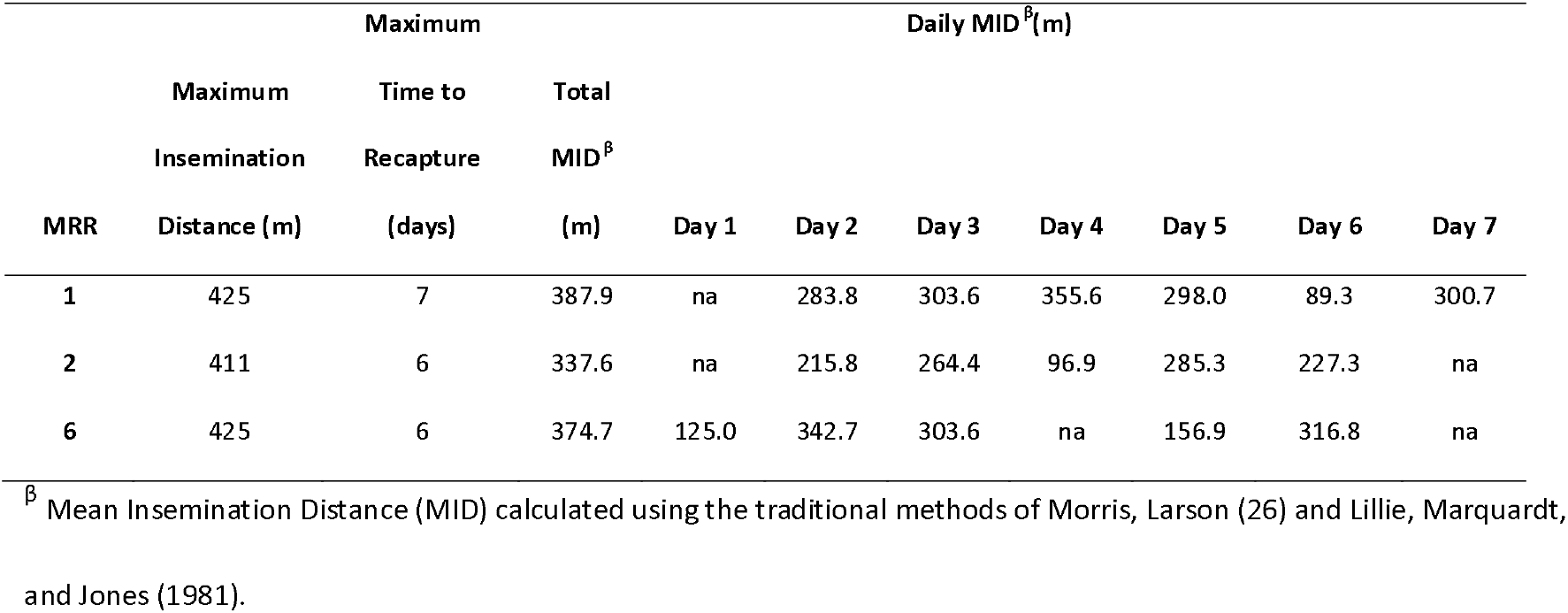
Mean Insemination Distances (MID) estimates adapted from traditional annulus-based methods. Estimates give an estimation of the distance at which inseminated females are then captured over time.

**Fig 7.**
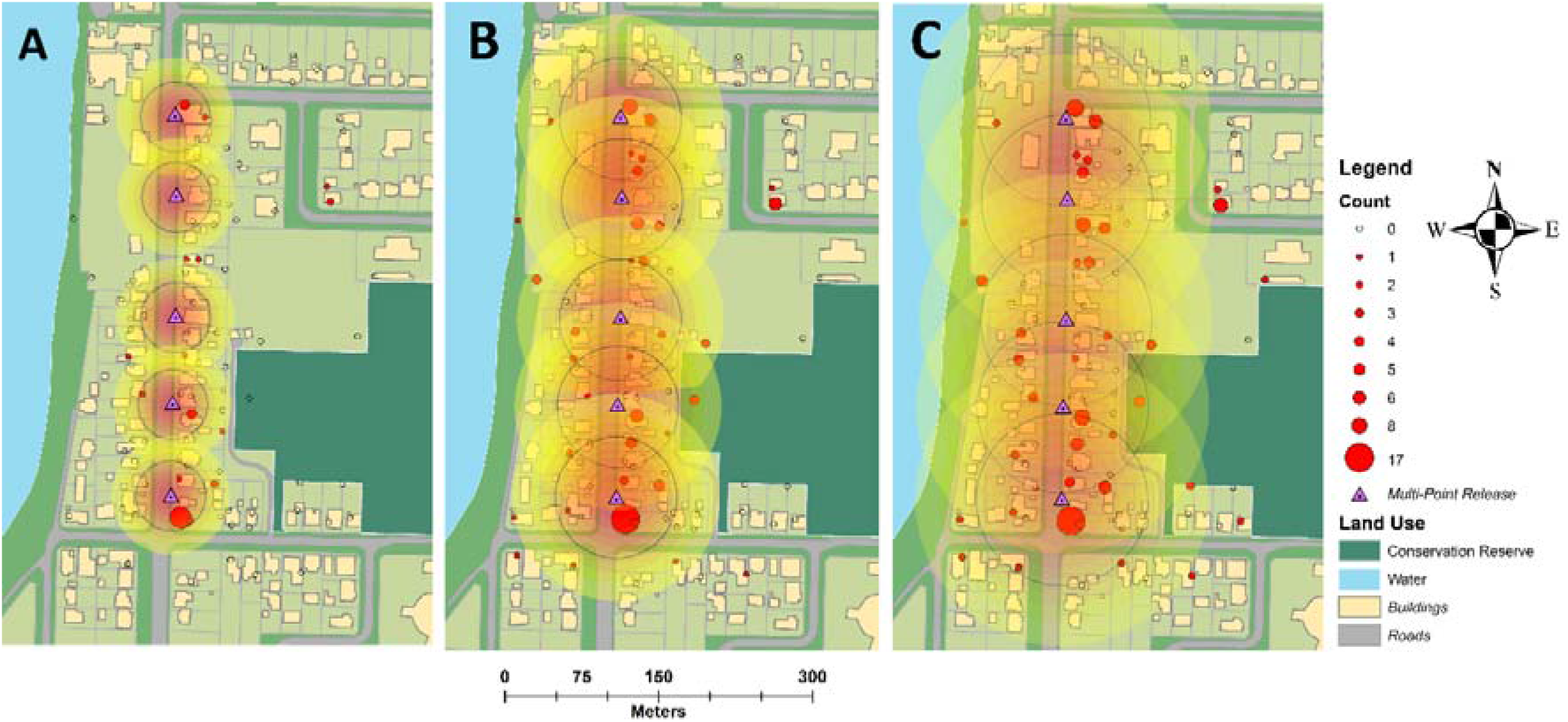
Multi-point estimates of *Aedes aegypti* movement during experiment four which apply the STEIK framework. Concentric circles from release point are density estimates of marked adult male *Aedes aegypti* one (A), three (B) and six (C) days post release. Black lines represent mean distance travelled over the time period.

### The role of urban landscape features

Contingency table analysis revealed Rhodamine B marked male *Ae. aegypti* were two times more likely to be captured in BGS traps situated at houses than in backyards or forests (χ^2^ = 10.06, df = 2, N = 564, Backyard OR = 0.47, 95 % CI = 0.28-0.77, Forest OR = 0.46, 95 % CI = 0.18-1.19, *P* < 0.007). Likewise, wild-type female *Ae. aegypti* had twice the likelihood of being captured in traps around houses than in backyards or forests (χ^2^ = 9.81, df = 2, N = 564, Backyard OR = 0.51, 95 % CI = 0.33-0.80, Forest OR = 0.57, 95 % CI = 0.24-1.37, *P* < 0.007). Wild-type males had a similar likelihood of being captured in traps around houses and forests (χ^2^ = 19.00, df = 2, N = 564, Forest OR = 1.09, 95 % CI = 0.50-2.49) but half the likelihood of being captured in backyard traps (OR = 0.47, 95 % CI = 0.32-0.69).

## Discussion

Vector control is entering a new era of area-wide, rear-and-release control strategies, where accurate measurements of population parameters will be essential for operational success (36, 41). Incompatible insect and sterile insect field interventions rely on several important biological parameters that determine the effectiveness of mass-reared male mosquitoes. Sterile insect interventions are reliant on the ability of mass-reared males to survive, seek out and successfully mate with available wild females. As female *Ae. aegypti* are typically thought to mate once during their lifetime, sterilization is reliant upon releasing enough mass-reared males (over-flooding) to outcompete wild males. Here we performed the first large-scale field releases of rhodamine B marked male *Ae. aegypti* into a wild population of mosquitoes. Our goal was to measure the basic biological parameters that govern movement, survival and mating interactions of mass-reared males within this population.

Competition between released and wild type male mosquitoes is difficult to discern within a heterogeneous population. Here, rhodamine B was used to mark the body and seminal fluid of male *Ae. aegypti*, enabling us to measure mating interactions with the wild female population and compare mating success between released and wild mates. Mating success of marked males was greatest when traps were not operating for the first day after release, suggesting that monitoring be reduced during the early phase of releases. Adding marked males into the wild population immediately impacted the proportion of inseminated females collected, however, this made no difference to the proportion of all inseminated females throughout each experiment (both rhodamine B and wild mated) and this was consistent across seasons. Furthermore, this trend was also constant in wild females inseminated by wild males across seasons. As release numbers were relatively similar across all experiments, these results suggest that the proportion of total mated females remains relatively static regardless of the number of additional males added to a population and sterile to wild-type mating ratios will be proportional to the number of each in the population.

This result is unlikely to be influenced by trapping bias (42). Wild females are likely available for mating immediately upon release of marked males, and that these females become available for mating at a relatively constant rate post-release, with ~40% of daily females captured at any given time being unmated. As *Ae. aegypti* males are limited by the rate at which they can mate during their lifetime (43), ensuring a constant supply of males into the landscape should be the priority of sterile male programs. Multiple releases of males will be required each week to ensure a constant and efficient overflooding ratio, a number that is highly reliant on the fitness of released males. Doing so will ensure a saturated landscape where females are mated by sterile males as soon as they become available and minimise the effect of immigration. If sterile males are not constantly present within the landscape, then programs will tend to be inefficient with higher costs and lower suppression than expected over time.

Observations during this study support the premise that male *Ae. aegypti* are capable of rapidly moving through a two-dimensional urban landscape (18). Studies on male *Ae. aegypti* movement tend to show large variations in distance travelled over the period of study and this has implications for how, where and when mosquito programs release males during SIT/IIT interventions. Results described here suggest male *Ae. aegypti* have the potential for long distance movement despite a short lifespan (17, 18). During single point releases marked males were recaptured at distances greater than 350 m and 400 m after day one and two, respectively. Furthermore, rhodamine B inseminated females were captured at the extent of the trapping array, with a high MID suggesting males have an effect greater than the area in which they are captured. Both isotropic and traditional MDT measurements of male movement between 95 m and 462 m (and across movement barriers) supports the general observation that the rate of male movement was greater than those traditionally recorded for females of the species. This would suggest that gene flow within an isolated population may be related to male spread as female movement is typically observed at less than 100 m over a lifetime (13, 44). This difference may be due to the different biological requirements of each sex, with females requiring a blood source, a resting place and oviposition site that may reduce dispersal, while the short life expectancy of males may have them constantly searching for virgin females. A greater understanding of dispersal patterns will result in maximum coverage across a landscape and lead to greater mating success, the primary goal of area-wide release programs.

While single point MRRs are ideal for measuring distance travelled over time, the utilization of measurements from multi-point releases are essential to optimizing area-wide releases. The STEIK estimates of movement used in this study (18, 39) benefited from the addition of temporal data and tended to produce larger estimates than those calculated through traditional methods. The STEIK method has the additional benefit of estimating movement without assuming individuals have been caught in traps. While greater male MDT was observed via single point than during multipoint releases, this is likely a function of both the release strategy (males being released across many traps) and a lack of clarity about which release sites captured mosquitoes had originated from (as a result of the single mark method employed). The latter point is important since our model considers that it was less likely that a mosquito caught close to one release point had travelled there from a distant release point, even though this could be the true movement pattern. During multi-point releases with a single marking type, the cumulative probability would therefore tend to be conservative when estimating MDT. As such, single point and multi-point releases cannot be easily compared when only a single marking type is used.

It is widely known that sterilization through transgenic and radiation approaches impose a fitness burden on released males (44, 45). Because of this, modern SIT programs have regularly failed due to lack of knowledge on sterile male performance post-release (41, 46, 47). Although we observed considerable differences in ALE between experiments, our results confirm the relatively short lifespan of male *Ae. aegypti* compared to that of females. It is estimated that female *Ae. aegypti* adults live on average for between five and nine days, with enough surviving the extrinsic incubation period to transmit pathogens during disease outbreaks (13). The differences between male and female lifespan has implications for scheduling the production and release of sterile male mosquitoes to achieve adequate overflooding of a wild population. If males used in a *Wolbachia* based IIT approach only survive for three to five days post-release, then releases will need to occur over regular intervals in order to ensure mass-reared males outcompete wild males and achieve the desired level of suppression. Release intervals will be presumably shorter for irradiated or transgenic males due to the additional fitness burden.

The LPI method is used historically for estimating population sizes during mark-release-recapture experiments. Ratios employed by this index compare the relationship between the total number of individuals captured, with the numbers released and then recaptured (8). Our results demonstrate the inadequacy of this method for estimating mosquito population sizes when recapture numbers are low. This finding is most evident when we compare population estimates (Table 2) with the total male and female captures for each experiment between seasons (Fig 2). The larger mean trapping rate of females during season two would suggest a population 1.5-2 times larger, however, the LPI estimates suggest otherwise. Although inaccuracies in the LPI are reflected by large confidence intervals, it is likely that even the lower CI estimates are unreflective of the population in the field, which when divided by the number of houses in the study area, suggest 31-57 males per house during season one. Previous results suggest this estimate is much higher than would be expected in the North Queensland region where the number of *Ae. aegypti* adults has been estimated as ca. 10 per residence during the wet season (48, 49). Furthermore, *Ae. aegypti* is an urban container inhabiting mosquito and populations are correlated with precipitation and temperature (48, 50). The lack of rainfall leading up to MRRs 1 and 2, when compared to the second season, would suggest a lower population leading up to and during the first season of experiments. Where the LPI provided the lowest variation and most likely most accurate population estimates (MRR 5 and 6), we recaptured a higher percentage of marked males when compared to wild males. Overall, results suggest that a well-mixed population is essential for accurate estimates of population size, and any estimates with low recapture rates should be considered with caution. Combining both the LPI and a measure of the insemination rate may lead to more accurate estimates of population size in the future experiments.

A lack of male *Ae. aegypti* movement studies has resulted in a deficiency of knowledge on individual behaviour and their ability to navigate through landscapes. This study placed traps in three major locations, house, backyard and forest, to observe how urban landscapes in north Queensland impact the movement of male *Ae. aegypti*. During both single and multi-point releases males tended to be recaptured within the same block they were released, and wind had little influence on the direction of movement. These findings support previous studies that suggest physical barriers influence movement between residential blocks (10, 18). However, roads or wind direction did not totally prevent male movement or mating with wild females in surrounding blocks, which suggests the open areas of Innisfail (such as roads or open grassy areas) only provided minimal resistance. Interestingly, both marked and unmarked males and females were captured in traps lining forest lines, with the majority of marked males in the first MRR caught in a forest trap directly behind a house at the single point release site. Forest traps also had the same likelihood of trapping wild males and females as traps situated in backyards. This observation supports previous studies that suggest that both male and female *Ae. aegypti* may instinctively move towards dark harbourage areas to seek shelter (51).

It is generally assumed that *Ae. aegypti* is most active during the daytime, as females show increased biting activity during the early morning and late afternoon (52). However, when males were released at night, we observed higher recaptures, a greater total MDT and a rapid mixing within the landscape when compared with daytime releases. It is possible that *Ae. aegypti* males move through the night, with low predation and movement boundaries (such as roads) playing less of a role than during daylight, thus increasing coverage across the landscape. Although we observed higher overall male movement during the night release, the lowest proportion of mated females during single-point releases was observed during this experiment. As the night release was not replicated caution should be assumed when interpreting these results and as such, additional studies are needed before firm conclusions can be drawn.

## Conclusion

The key to the next era of rear and release vector control will lie in the capacity of authorities to release competitive males that disperse widely, survive for greater periods and interact effectively with wild females. While the scientific literature contains extensive detail on female movement, there is relatively little quantitative information detailing the behaviour of wild-type male *Ae. aegypti*, and no studies exploring insemination rates in a field setting. Not only does rhodamine B provide new insights into male *Ae. aegypti* movement characteristics in urban landscapes, but additional information on how efficacious mass released male mosquitoes are at searching for and inseminating females in a wild population. The unique insights provided by our study into male *Ae. aegypti* biology will lay a foundation for designing and optimizing robust and effective male release strategies in the future and lead to a greater understanding of mating interactions in the wild.

## Supporting information

Supplemental Images 2

## Acknowledgments

We would like to thank Matt Bradford, Caleb Anning and Ben Purcell for assistance in trap placement, public engagement and mosquito collections. Also, to Helen Cook for her contribution and fearless work ethic in ensuring the community remained successfully engaged during the extensive study period. We would like to acknowledge Kamran Najeebullah at CSIRO for wind direction plots relative to mosquito recaptures. Finally, we would like to thank the Cassowary Coast community around Innisfail for their kind support and hosting our mosquito traps throughout the research.

## Supporting Information

**S1 Table. Rainfall during, two weeks before, mean daily minimum and maximum temperatures and mean relative humidity during the experimental MRR periods.**

**S2 Figure. Primary wind directions during MRR experimental periods.**

**S3 Figure. Daily recaptured mosquitoes relative to wind direction during experimental periods.**

## Data reporting

Stored male recapture data and R code for the STEIK framework is available at https://github.com/dpagendam/MRRk. Raw mosquito and trap data is available at https://data.csiro.au/collections/collection/CIcsiro:47209v1#/collection/CIcsiro:47209

## Financial Disclosure Statement

This work was supported by the Australian National Health and Medical Research Council (NHMRC 1082127). Project partners at Verily Life Sciences provided assistance in study design and reviewed the final manuscript but played no role in data collection, analysis or the decision to publish.

## Competing interests

There are no competing interests, whether they be financial, personal or professional, that have influenced the work.

## Human and animal research

Human ethics was sought through the CSIRO Social and Interdisciplinary Science Human Research and Ethics Committee (CSSHREC) and approved under project 026/16 named “Sterile insect technology development for *Aedes aegypti*”.

---

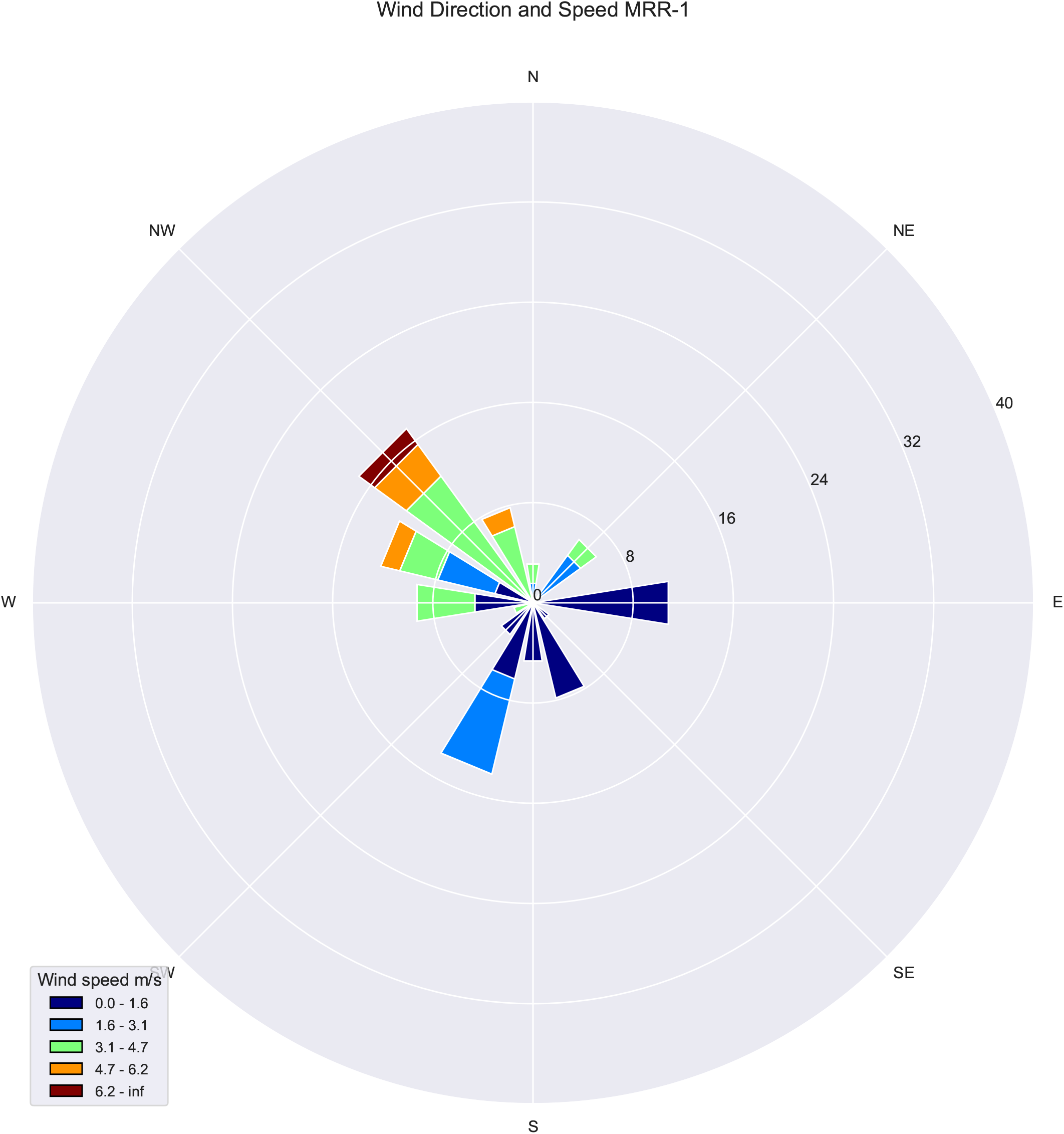

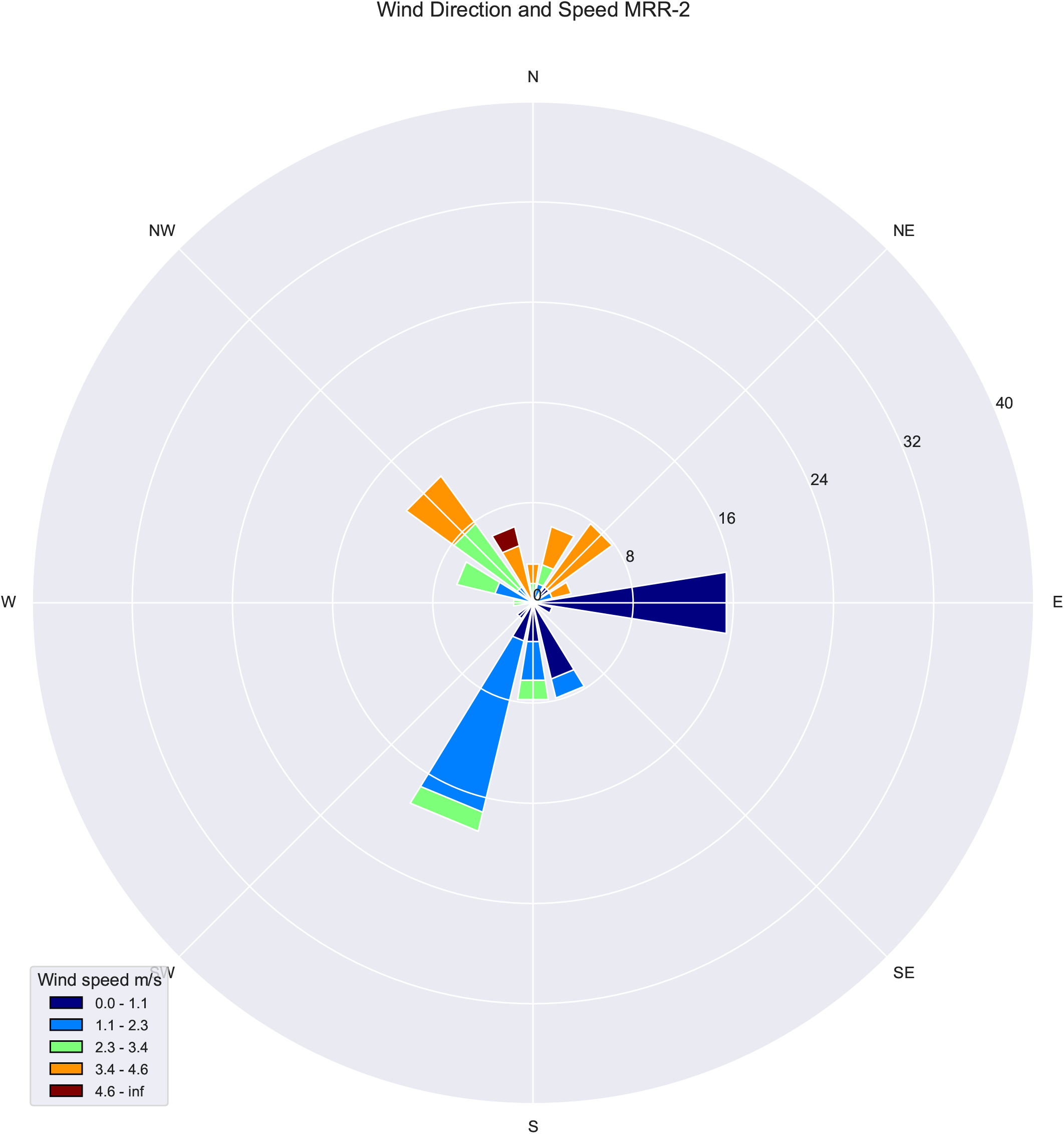

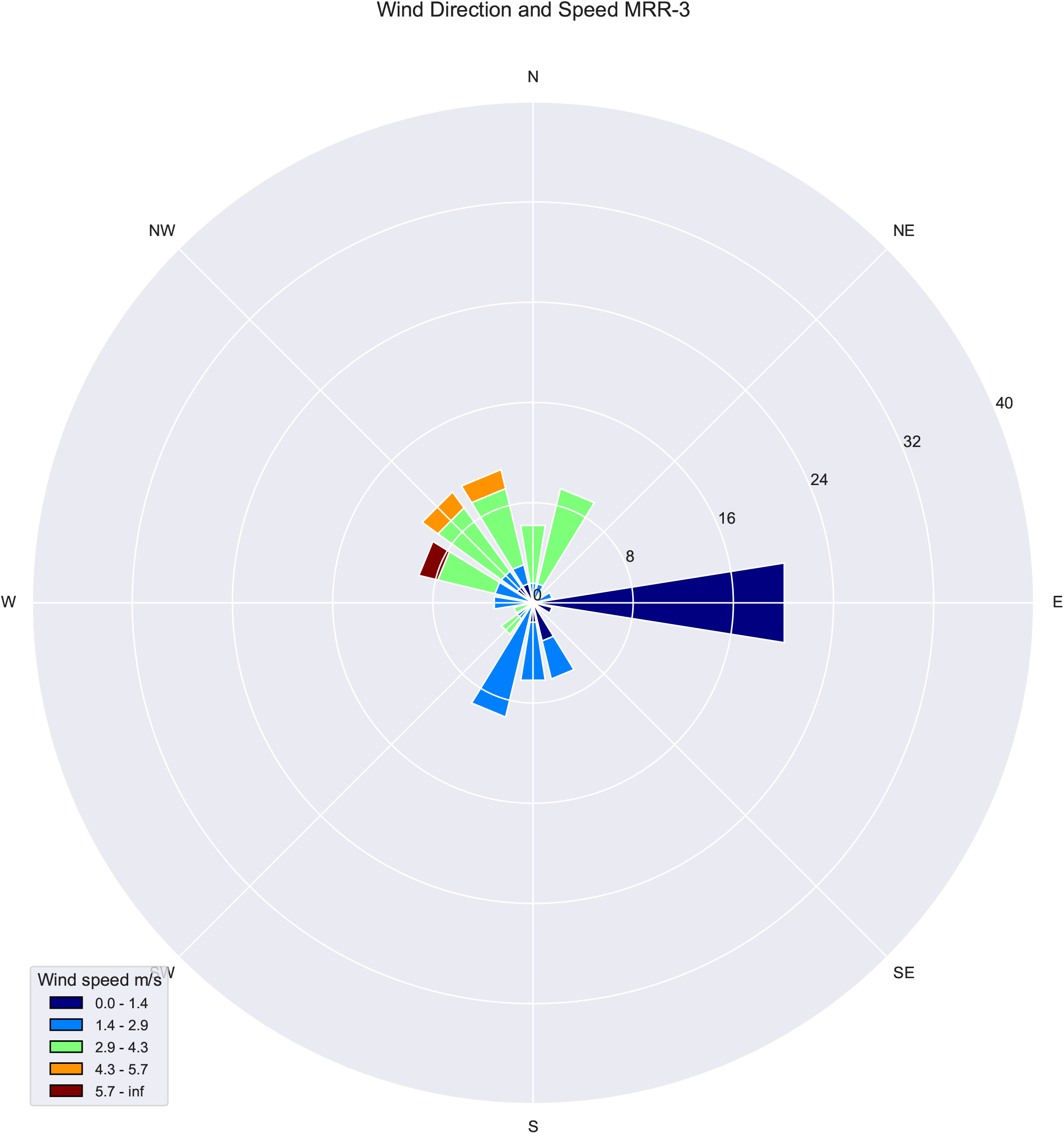

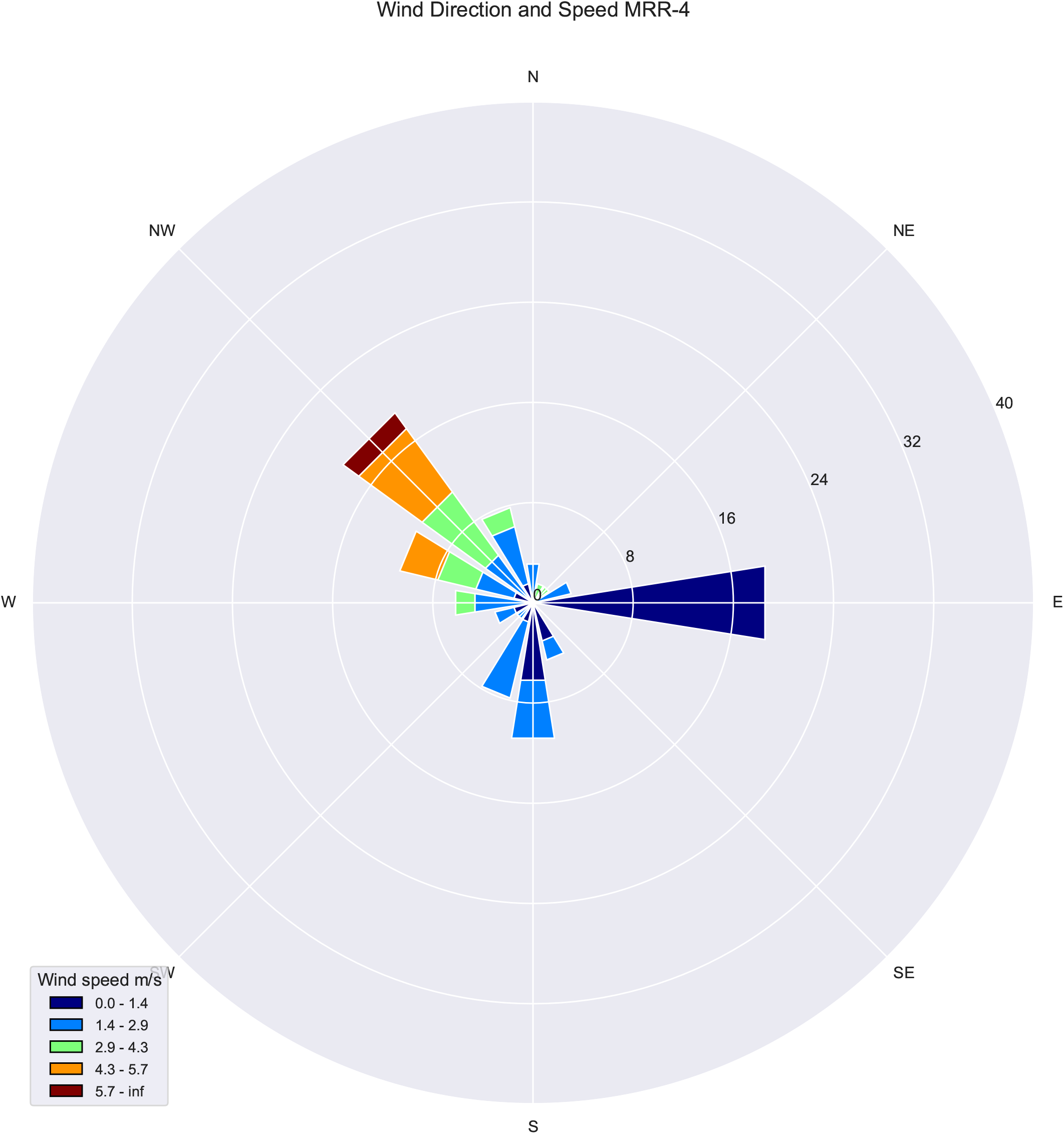

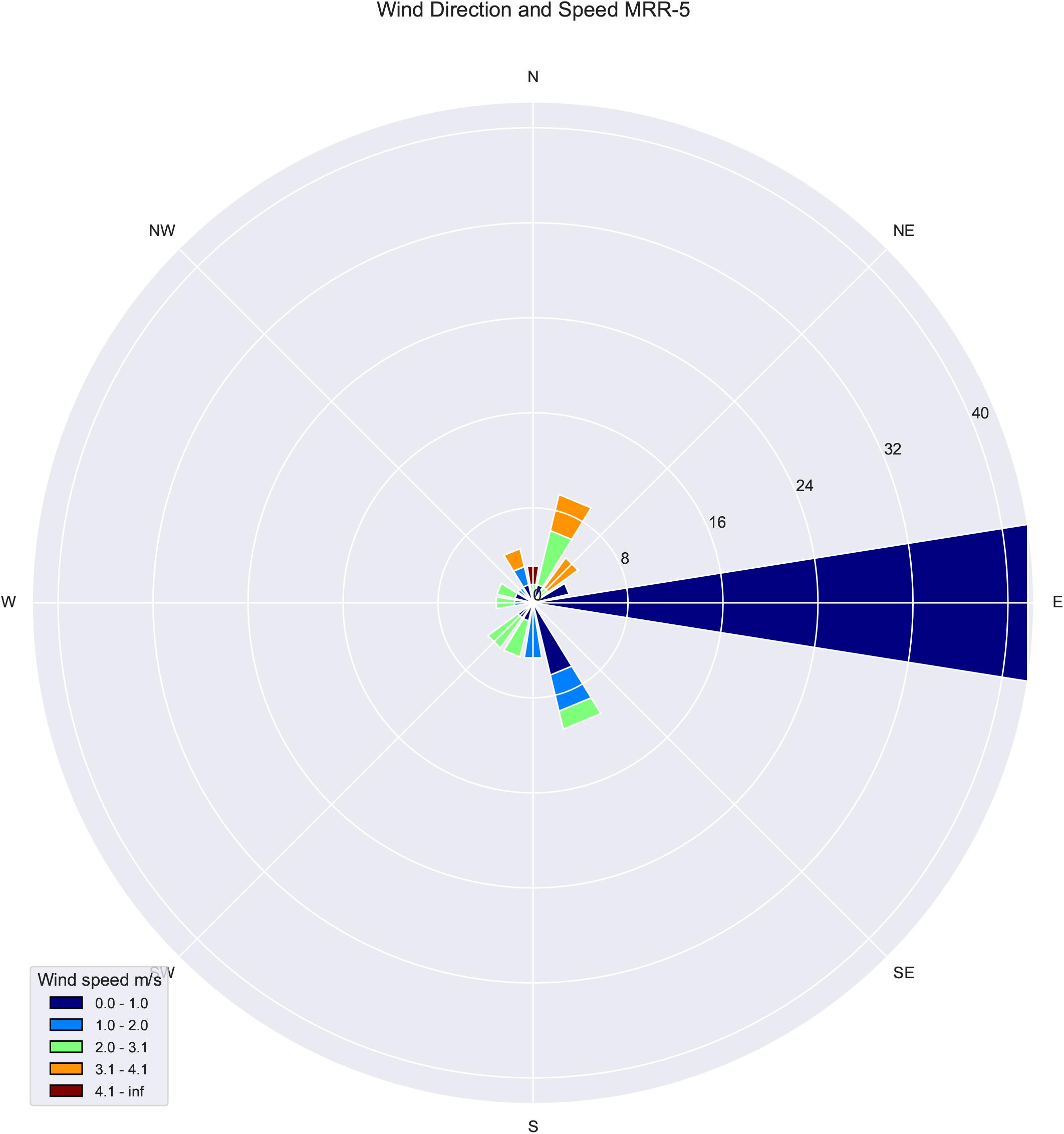

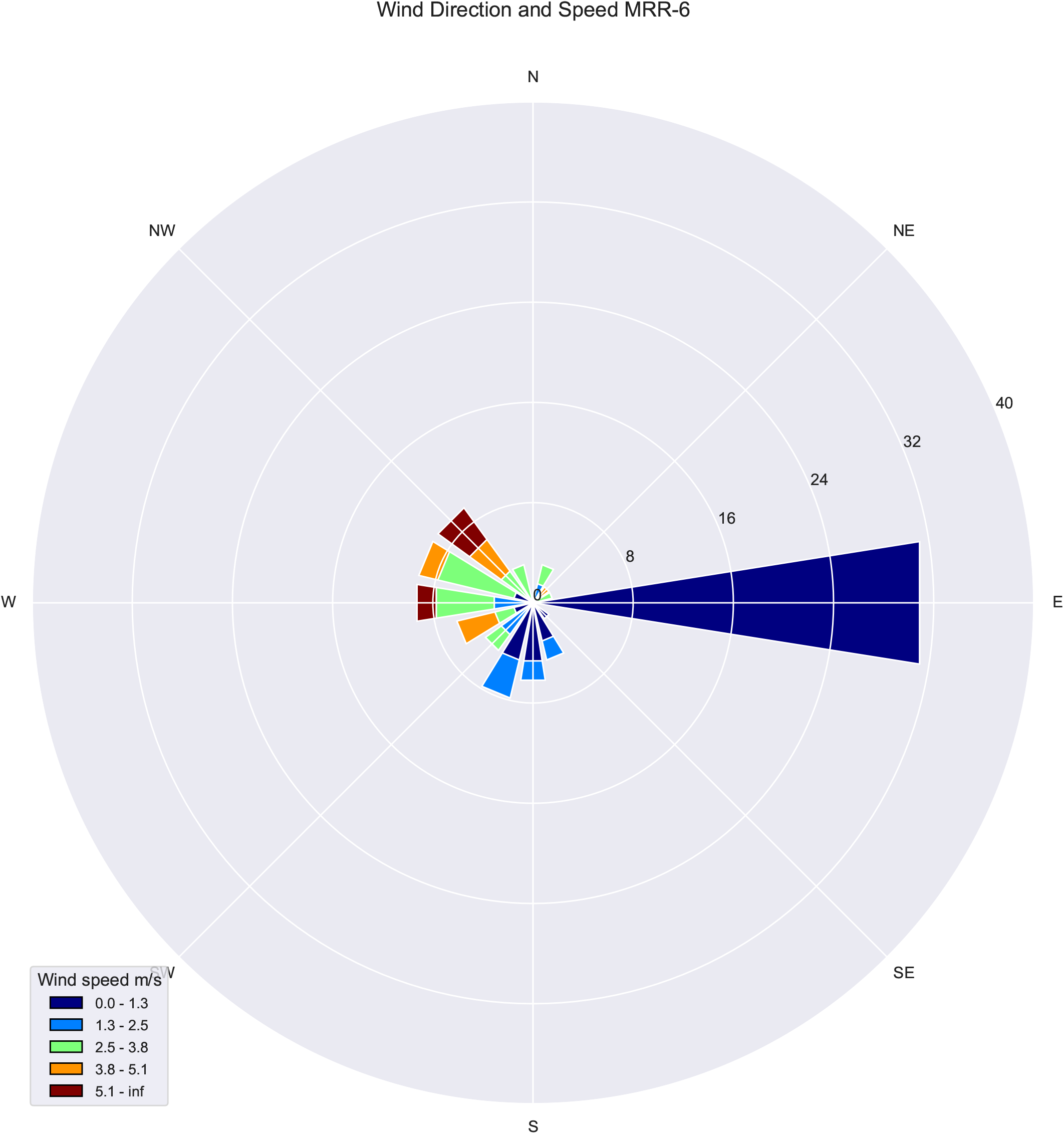

## Notes

https://doi.org/10.25919/5f9f642323d86

https://github.com/dpagendam/MRRk

### Competing Interest Statement

The authors have declared no competing interest.

