## Supplemental Table 1 for "Mark-release-recapture of male *Aedes aegypti* (Diptera: Culicidae): use of rhodamine B to estimate movement, mating and population parameters in preparation for an incompatible male program"

**S1 Table. Rainfall during, two weeks previous to, mean daily minimum and maximum temperatures and mean relative humidity during the experimental MRR periods.**

| **MRR** | **Mean Minimum Temperature (SD)** | **Mean Maximum Temperature (SD)** | **Total**  **Rainfall During Experiment (mm)** | **Total Rainfall 2 Weeks Previously (mm)** | **Mean Humidity**  **9am**  **(SD)** | **Mean Humidity 3pm**  **(SD)** |
| --- | --- | --- | --- | --- | --- | --- |
| **1** | 20.8 (0.9) | 30.2 (1.0) | 39 | 14 | 68.8 (11.3) | 56.9 (6.4) |
| **2** | 21.0 (1.4) | 31.6 (0.8) | 0 | 51 | 63.3 (4.1) | 53.8 (6.8) |
| **3** | 22.0 (1.1) | 31.1 (1.6) | 168 | 0.8 | 68.0 (16.3) | 65.0 (17.3) |
| **4** | 23.3 (0.8) | 31.7 (1.6) | 151 | 242 | 78.4 (9.2) | 69.6 (8.7) |
| **5** | 23.8 (1.8) | 32.7 (2.9) | 95 | 344 | 63.0 (26.9) | 72.1 (14.3) |
| **6** | 23.4 (0.7) | 30.8 (0.8) | 154 | 176 | 81.1 (8.8) | 73.1 (9.9) |
