## Supplemental Images 2 for "Mark-release-recapture of male *Aedes aegypti* (Diptera: Culicidae): use of rhodamine B to estimate movement, mating and population parameters in preparation for an incompatible male program"

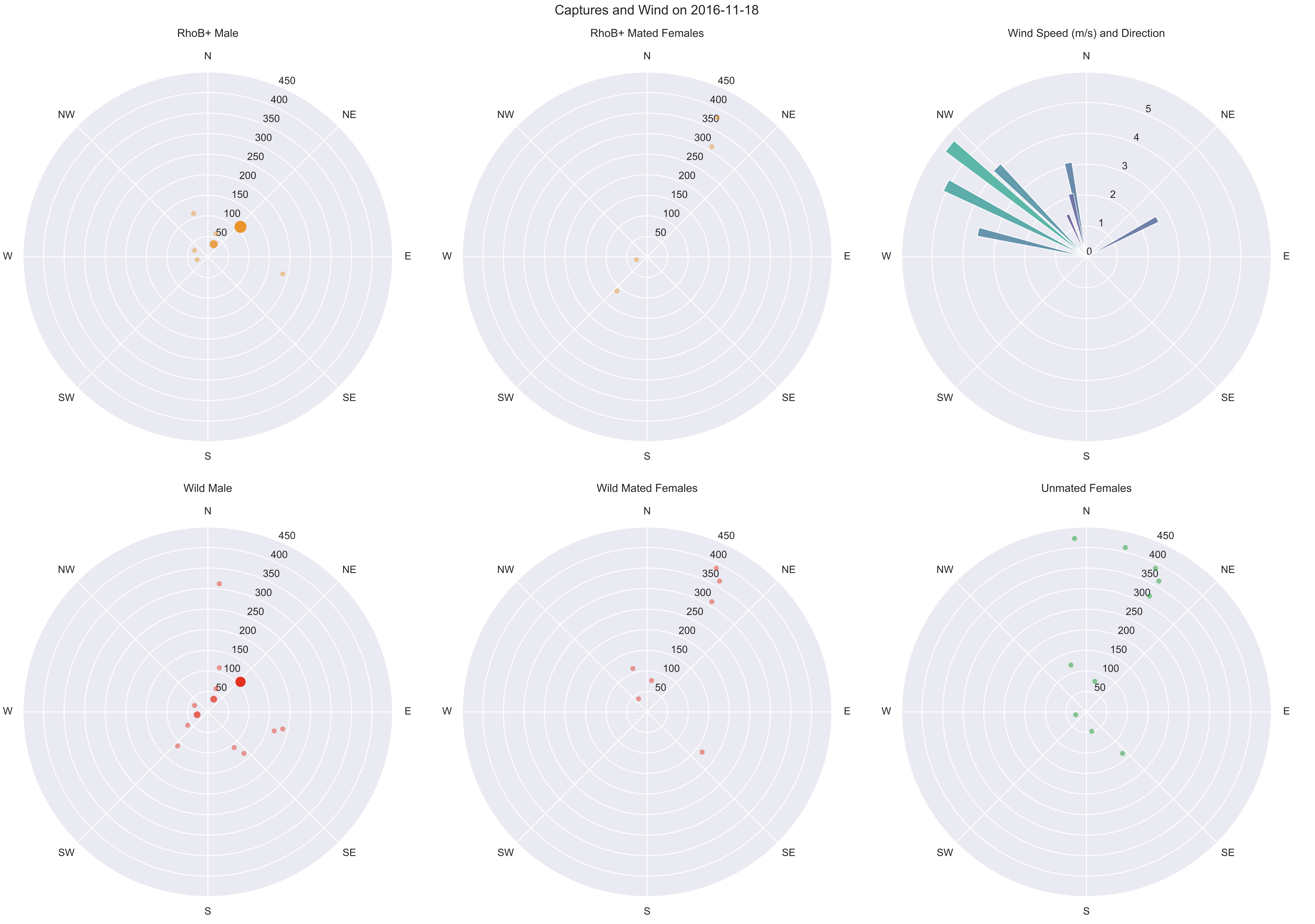
Captures and Wind on 2016-11-18
RhoB+ Male
RhoB+ Mated Females
Wind Speed (m/s) and Direction
Wild Male
Wild Mated Females
Unmated Females
N
NE
E
SE
S
SW
W
NW
50
100
150
200
250
300
350
400
450
0
1
2
3
4
5

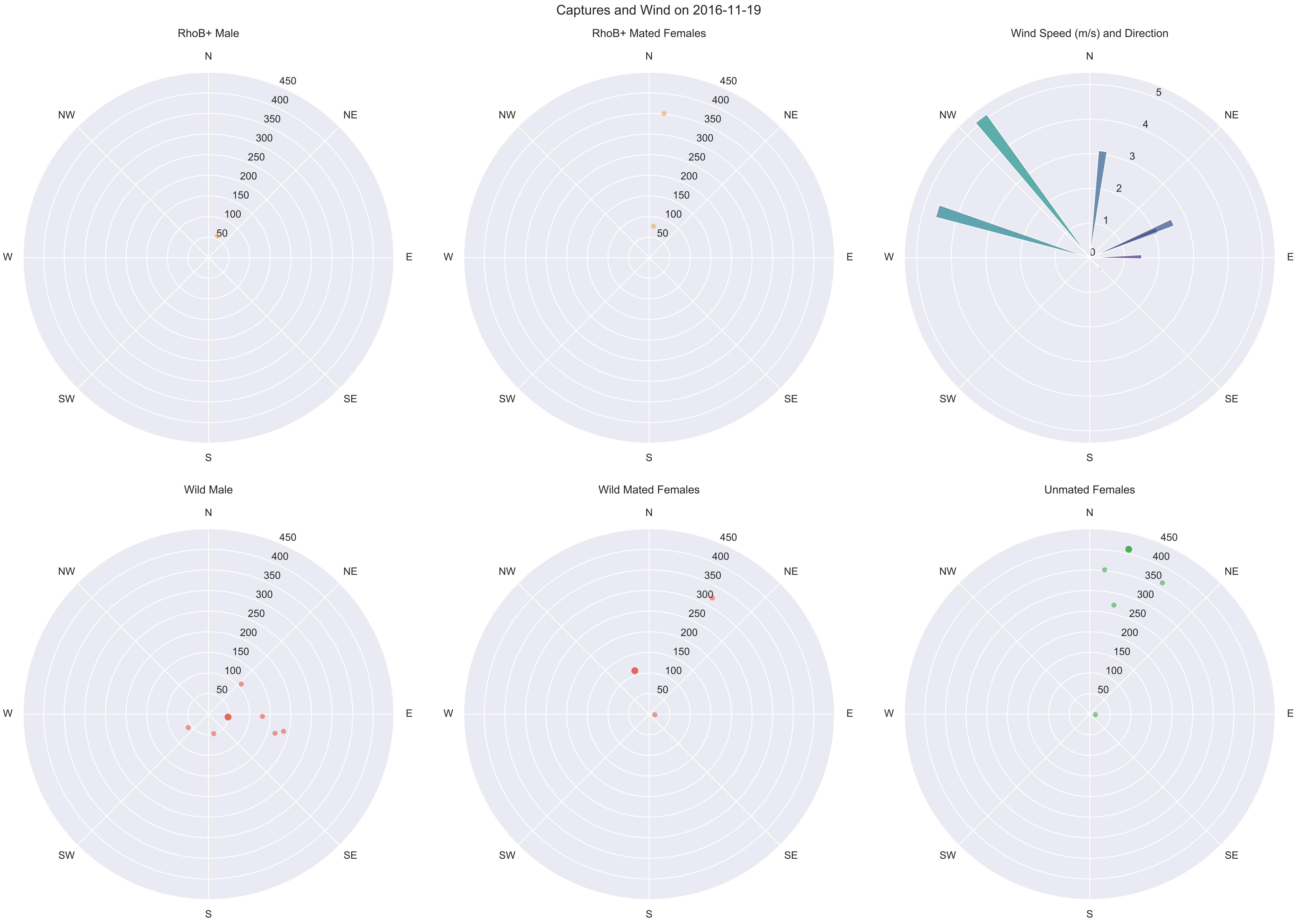

Captures and Wind on 2016-11-19
RhoB+ Male
RhoB+ Mated Females
Wind Speed (m/s) and Direction
Wild Male
Wild Mated Females
Unmated Females
N
NE
E
SE
S
SW
W
NW
50
100
150
200
250
300
350
400
450
0
1
2
3
4
5

Captures and Wind on 2016-11-20

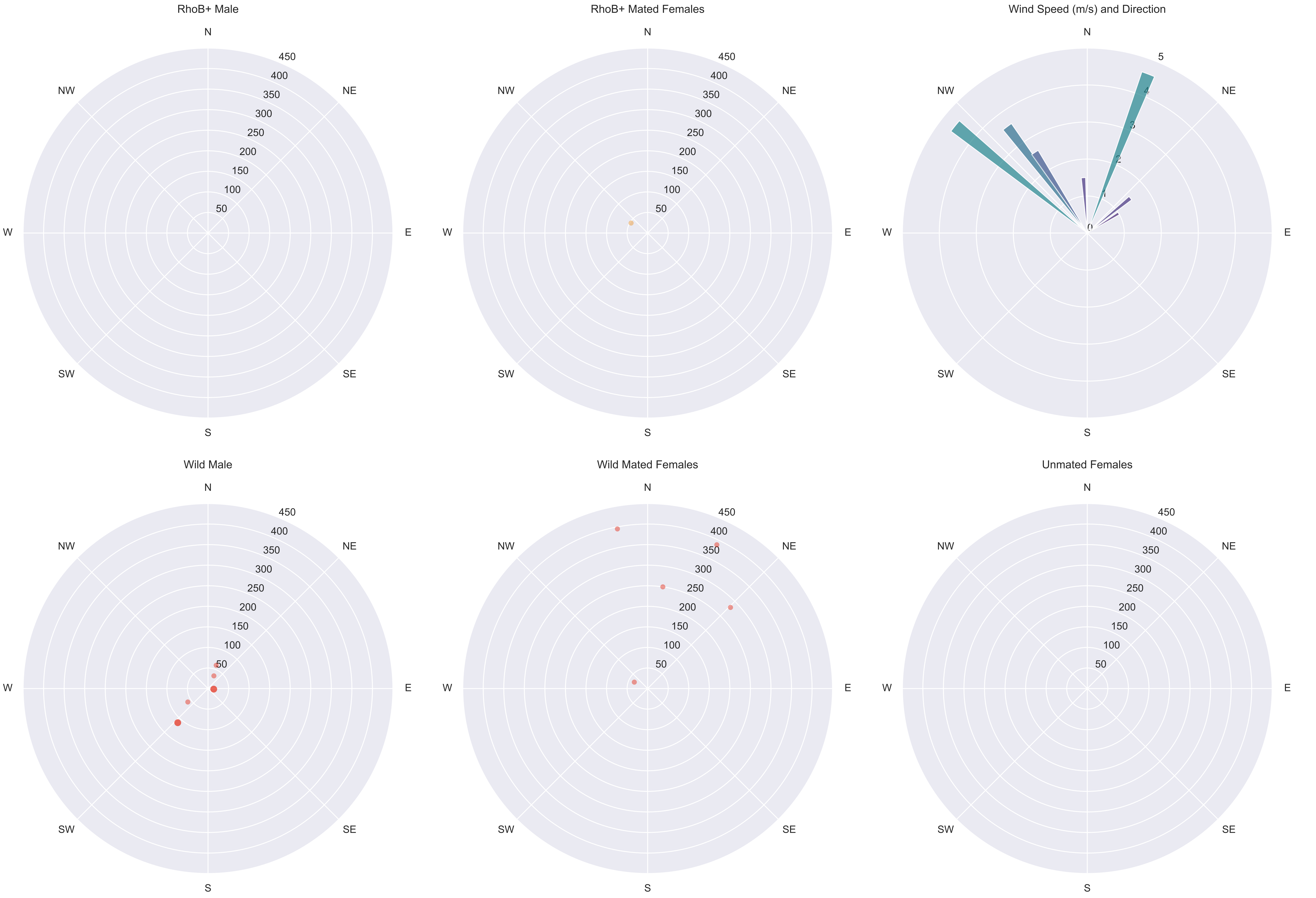

Captures and Wind on 2016-11-21

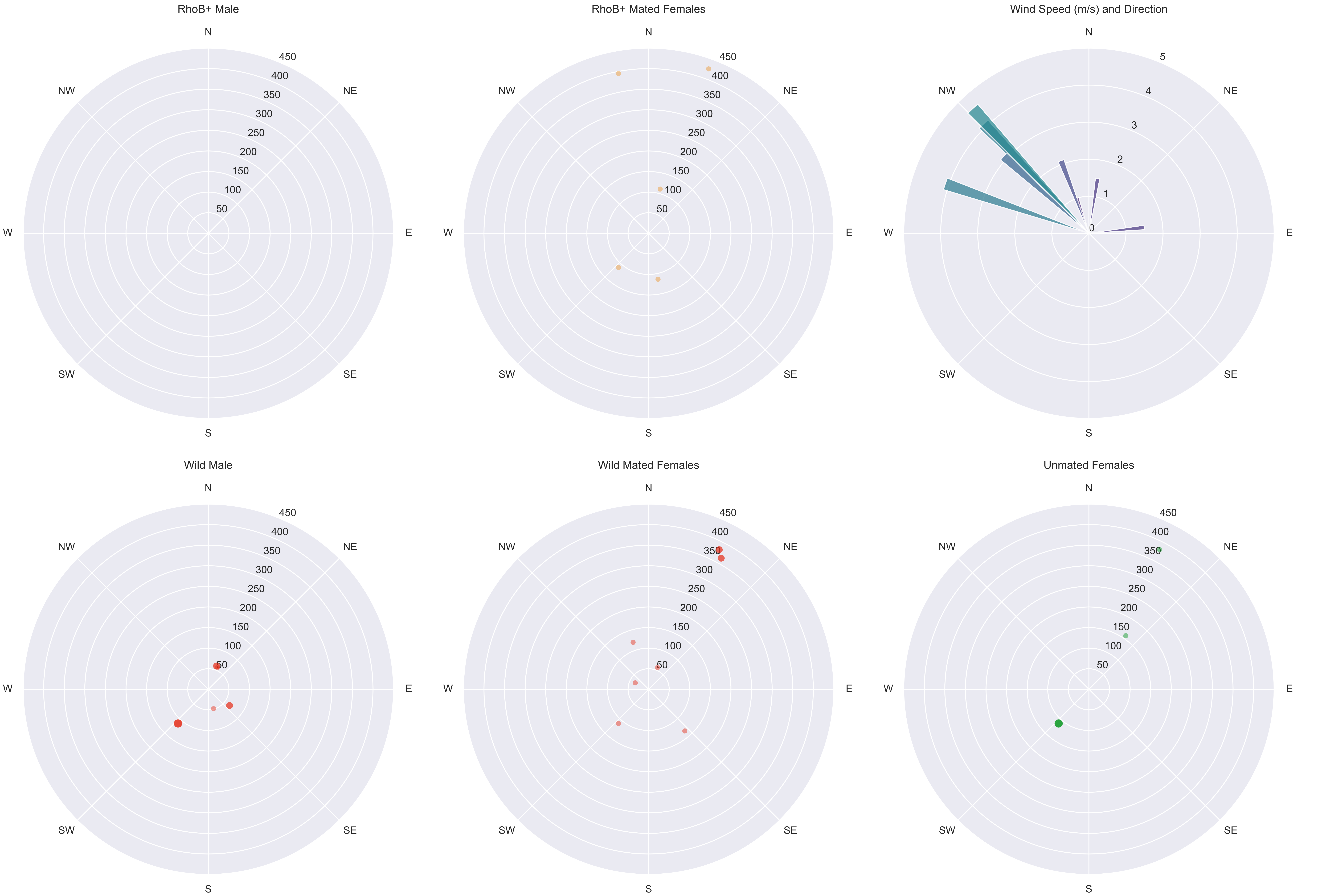

Captures and Wind on 2016-11-22

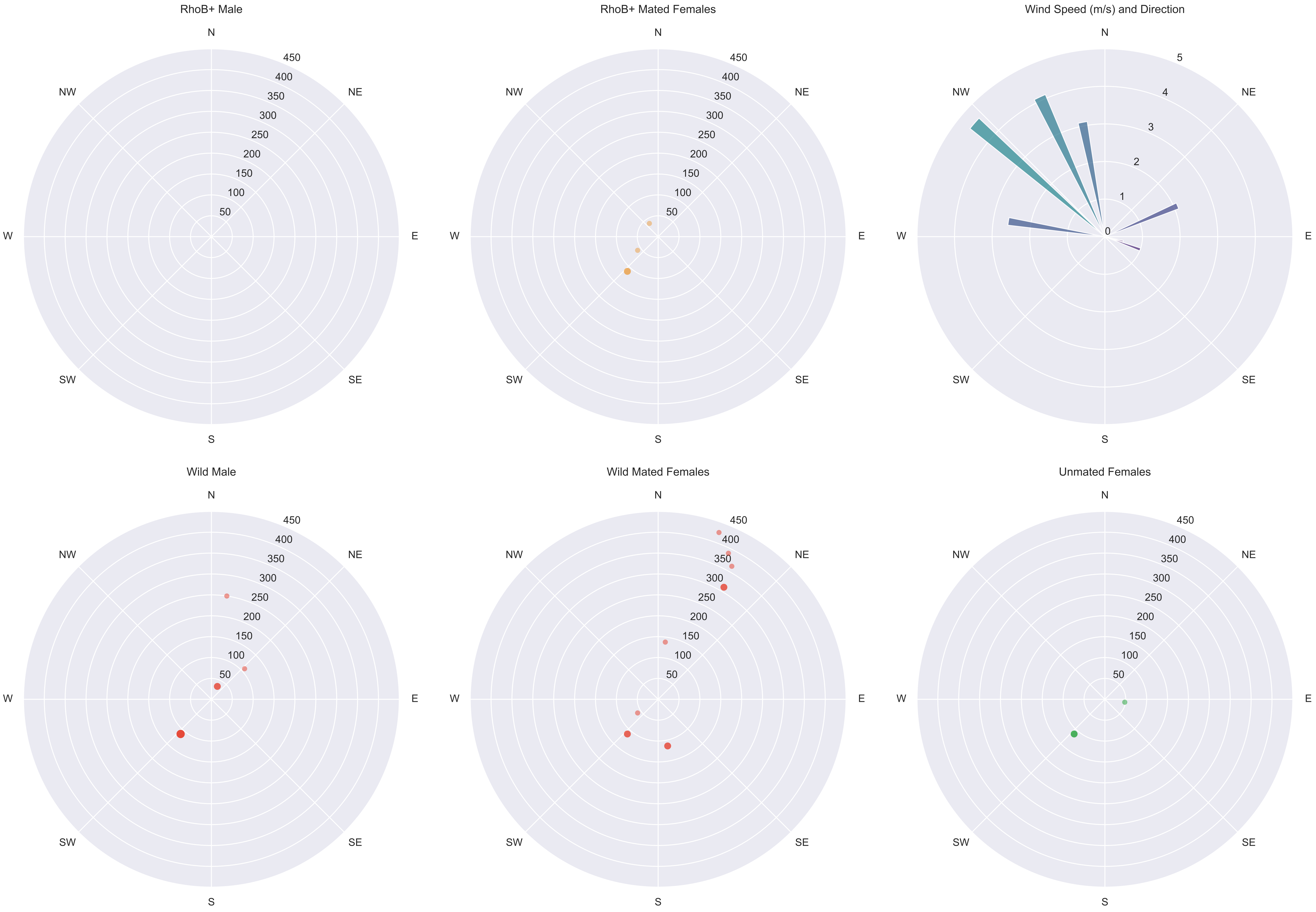

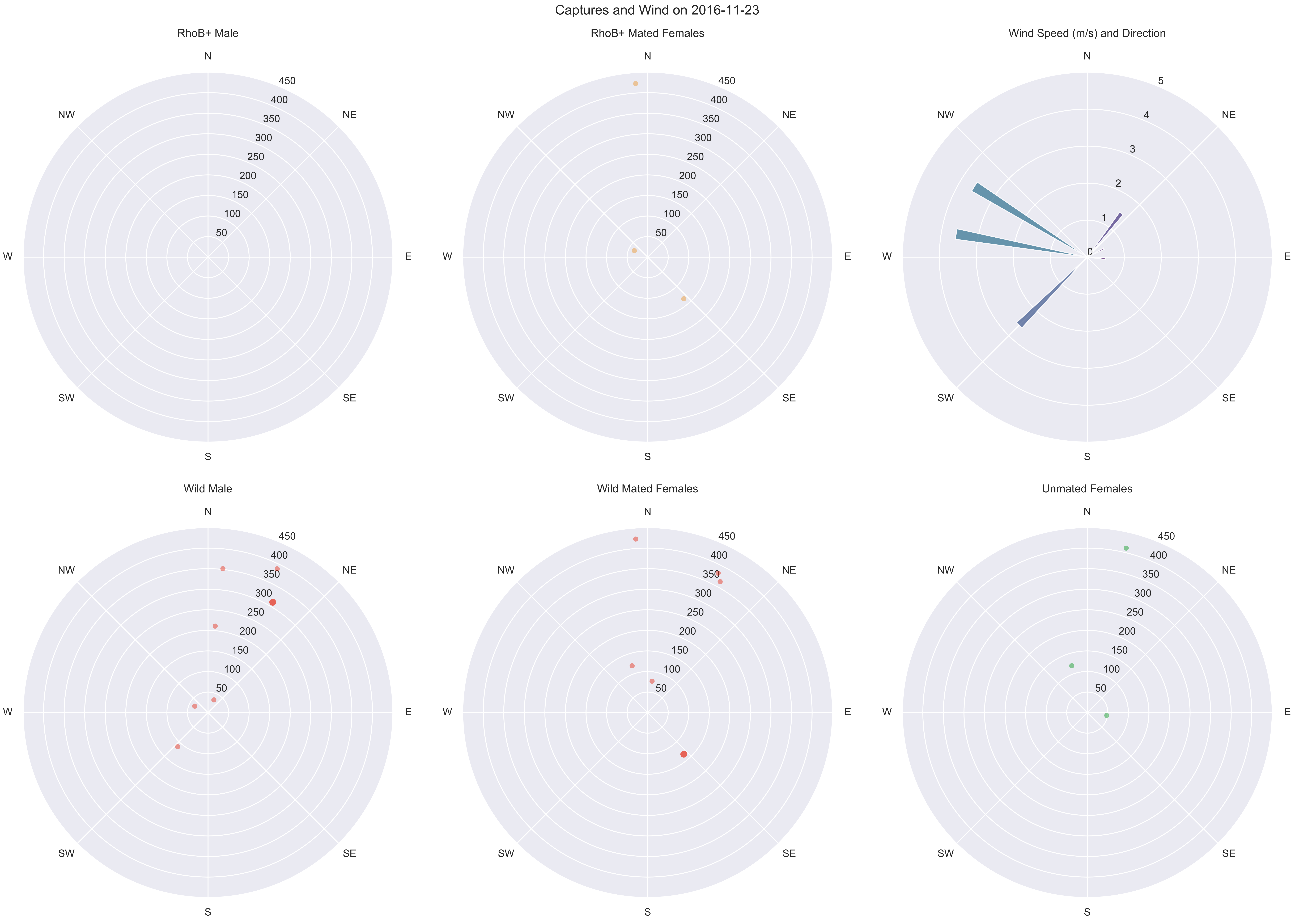

Captures and Wind on 2016-11-23
RhoB+ Male
RhoB+ Mated Females
Wind Speed (m/s) and Direction
Wild Male
Wild Mated Females
Unmated Females
N
NE
E
SE
S
SW
W
NW
50
100
150
200
250
300
350
400
450
0
1
2
3
4
5

Captures and Wind on 2016-11-24

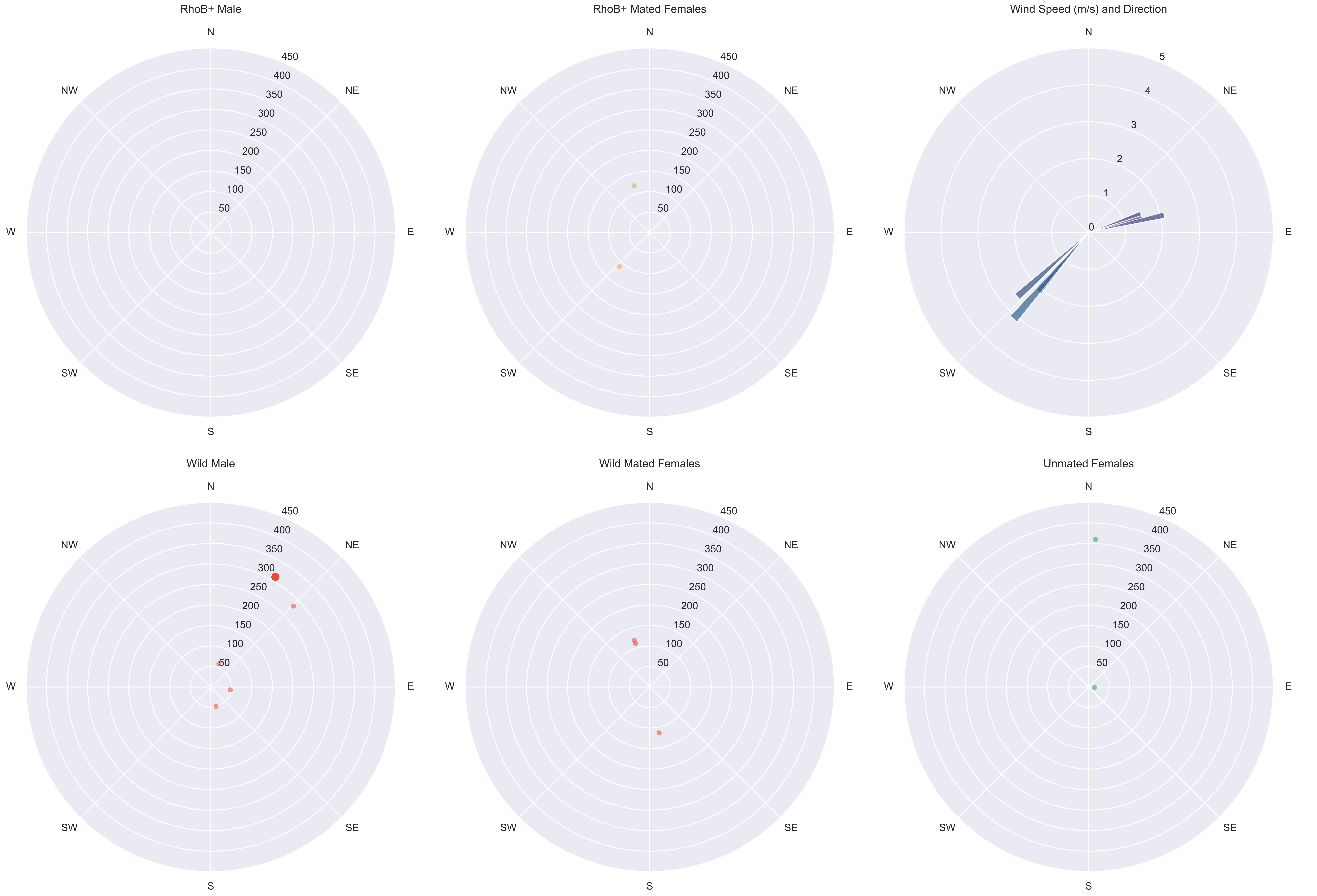

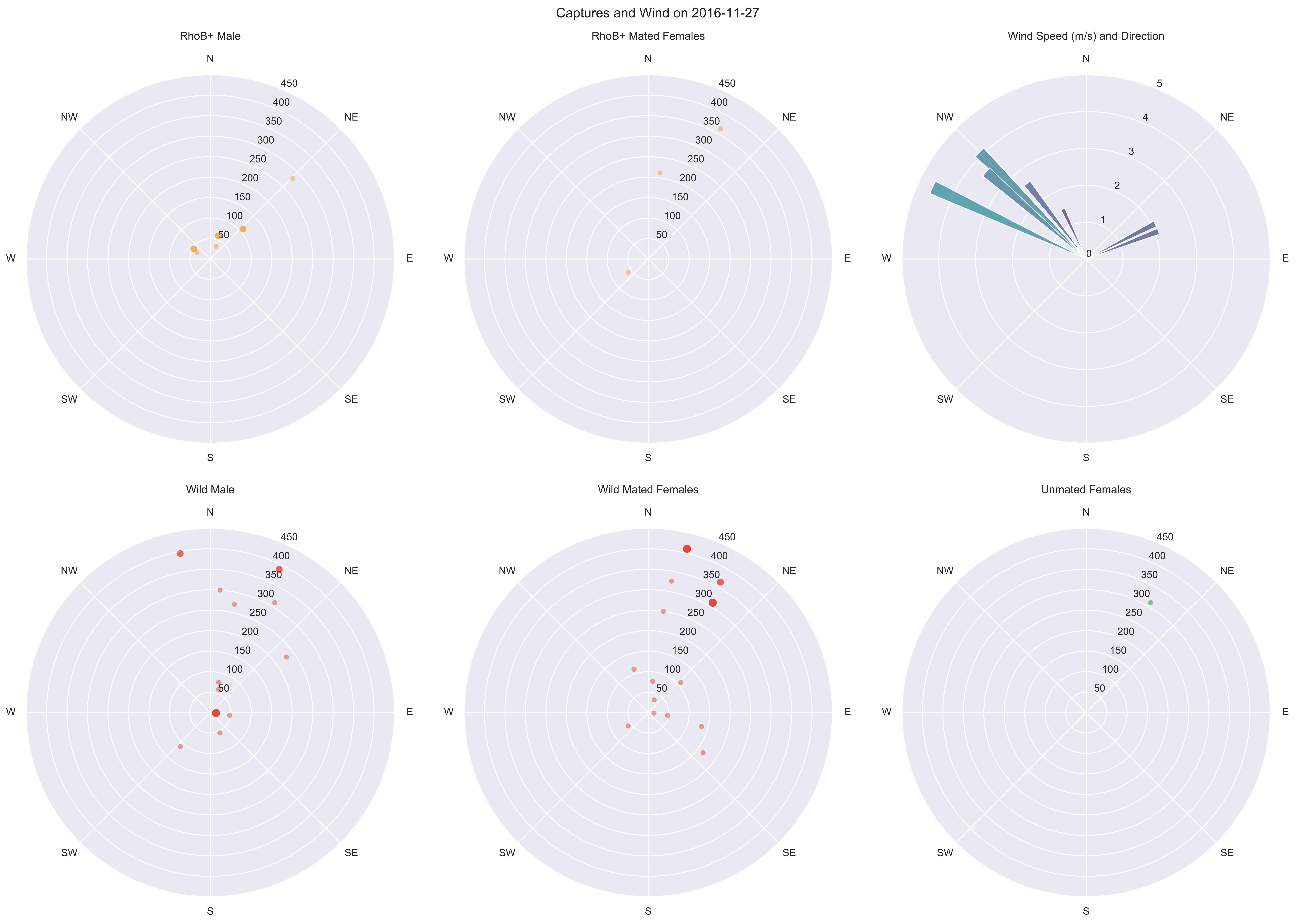
Captures and Wind on 2016-11-27
RhoB+ Male
RhoB+ Mated Females
Wind Speed (m/s) and Direction
Wild Male
Wild Mated Females
Unmated Females
N
NE
E
SE
S
SW
W
NW
50
100
150
200
250
300
350
400
450
0
1
2
3
4
5

Captures and Wind on 2016-11-28

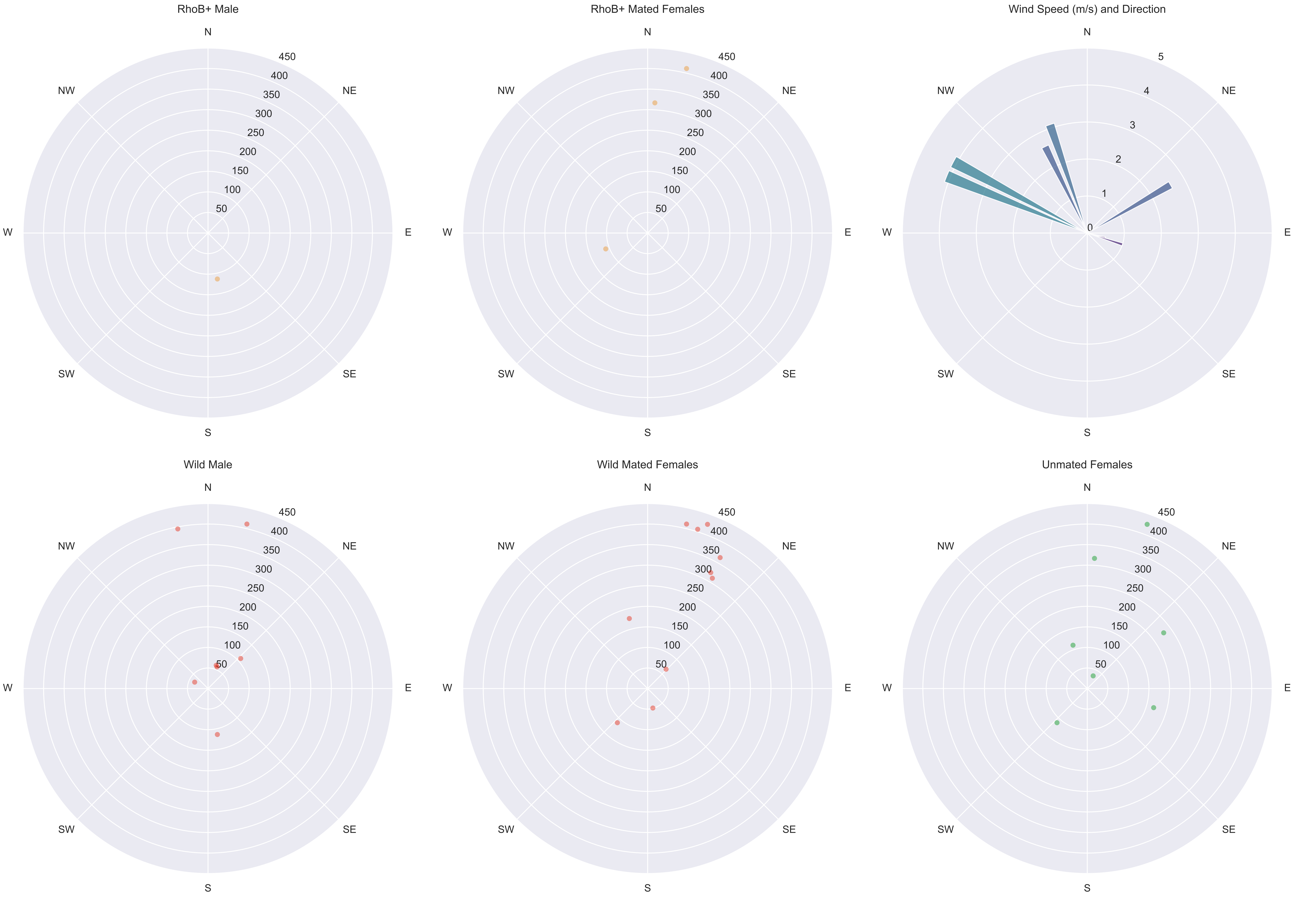

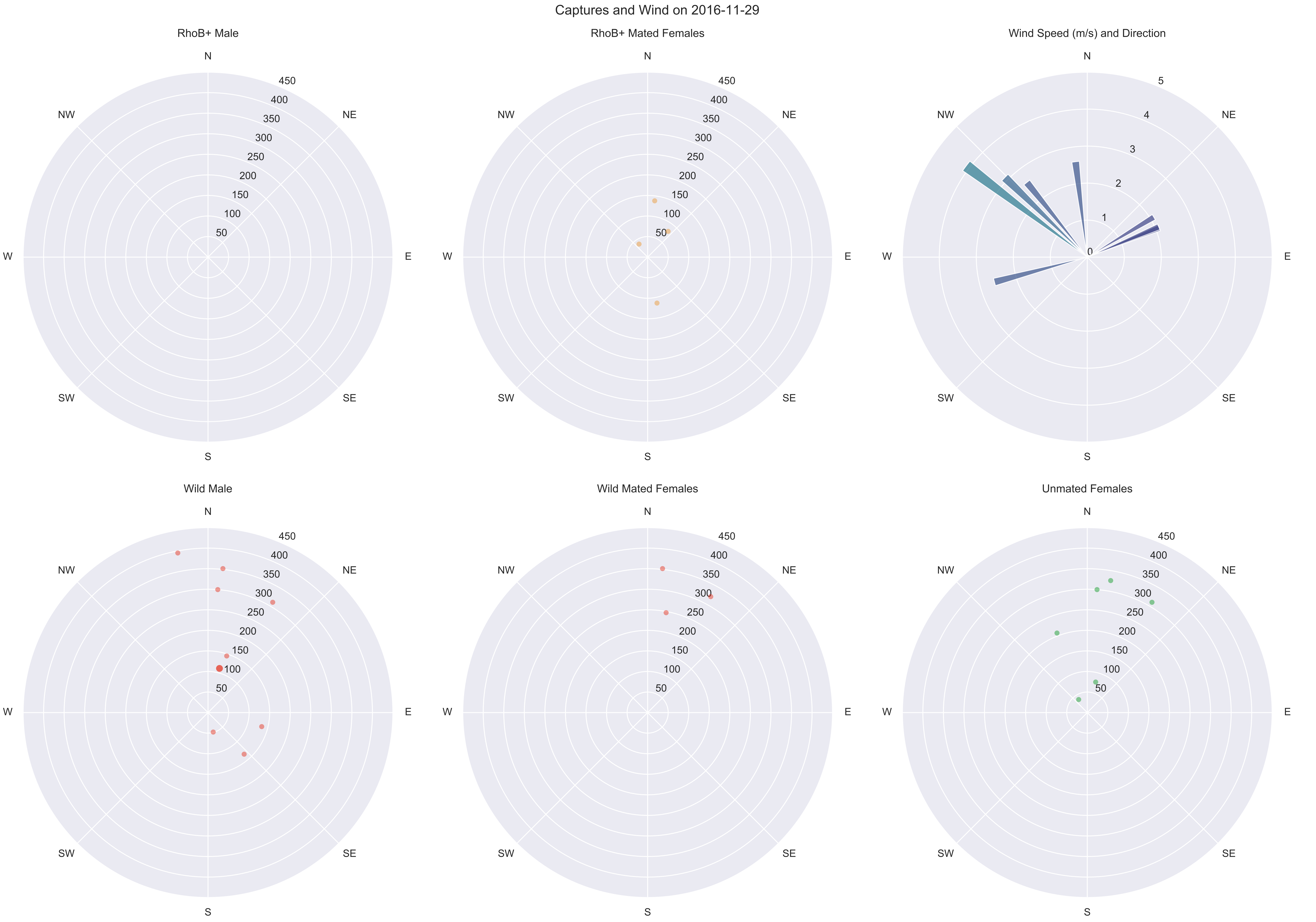

Captures and Wind on 2016-11-29
RhoB+ Male
RhoB+ Mated Females
Wind Speed (m/s) and Direction
Wild Male
Wild Mated Females
Unmated Females
N
NE
E
SE
S
SW
W
NW
50
100
150
200
250
300
350
400
450
0
1
2
3
4
5

Captures and Wind on 2016-11-30

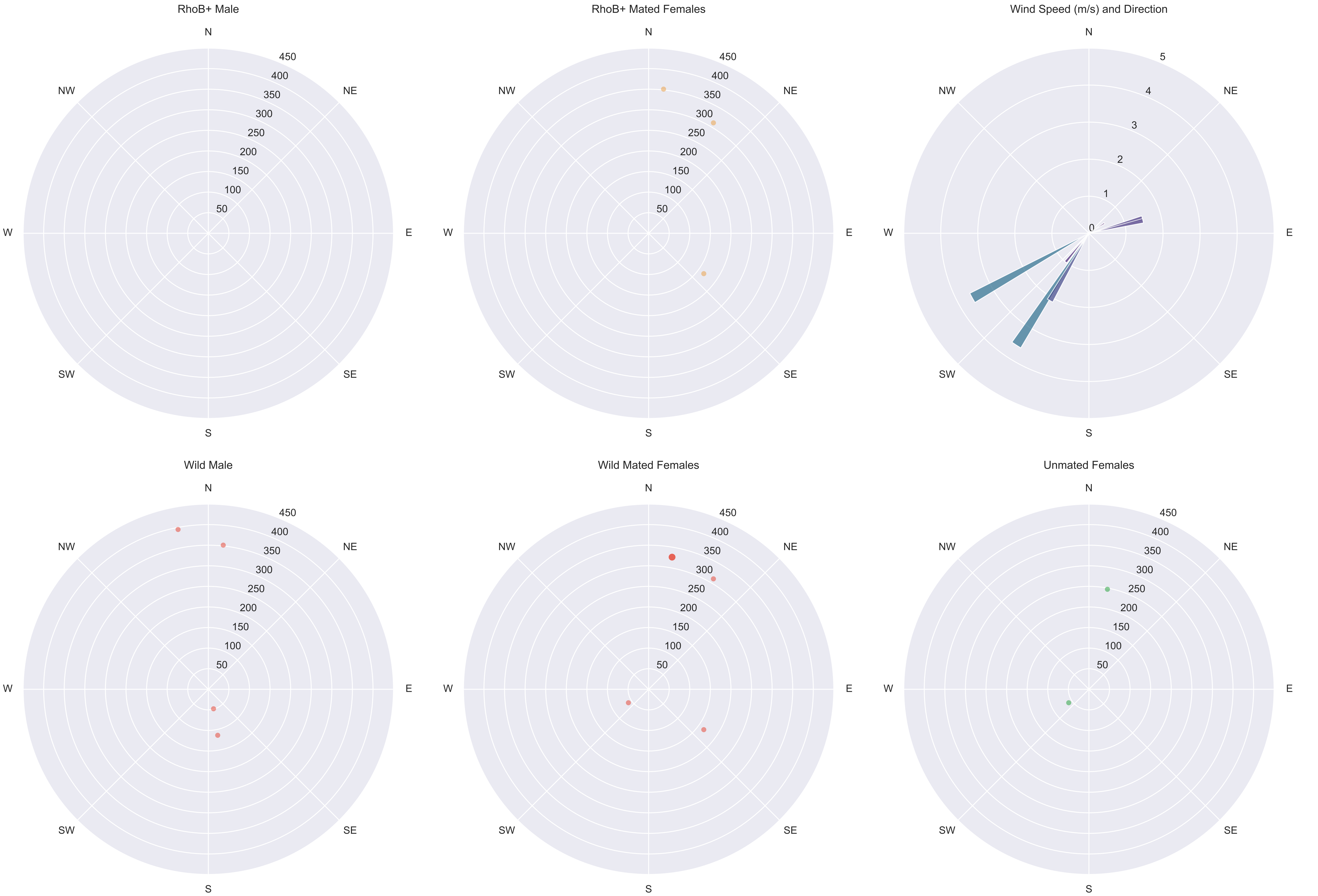

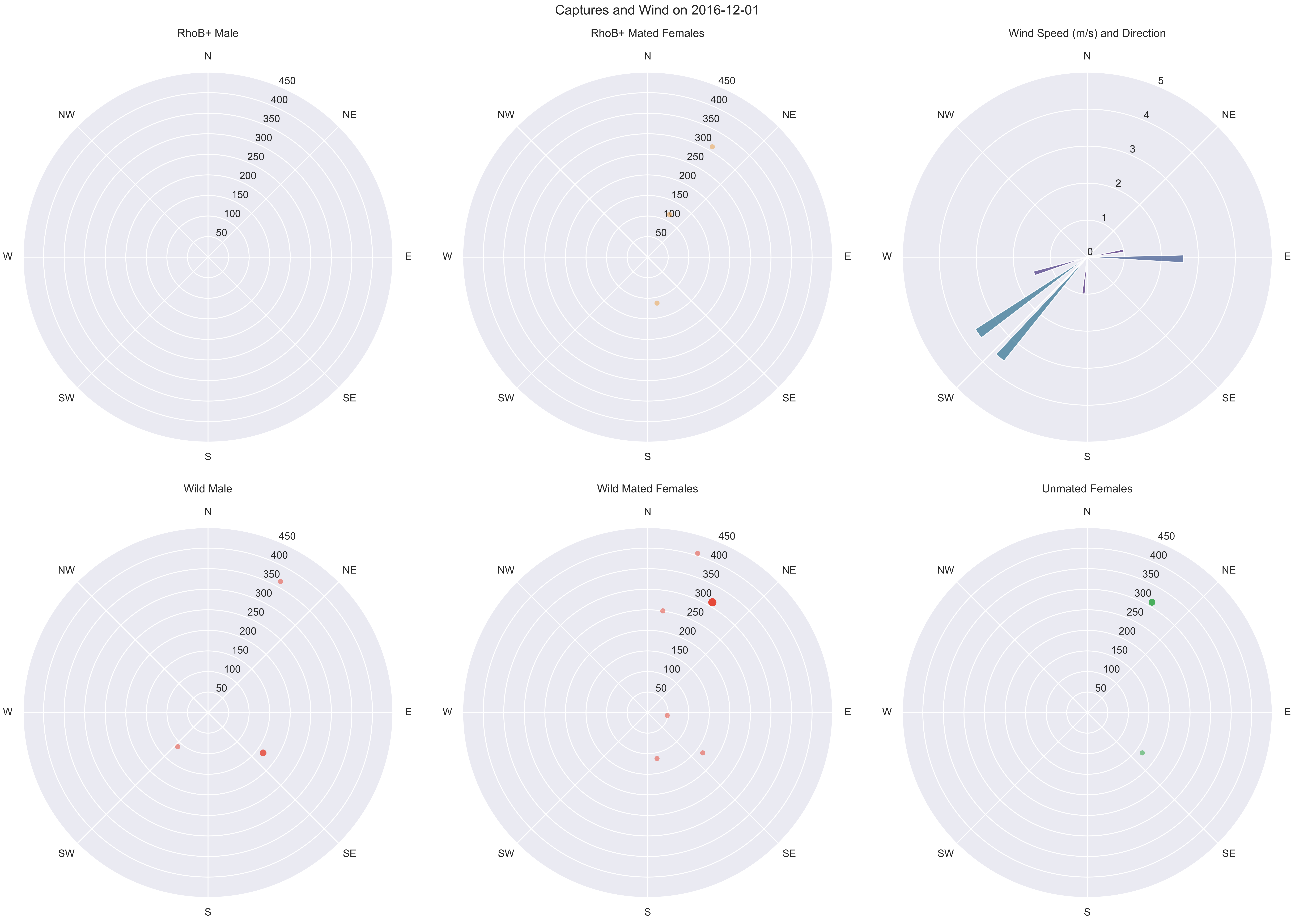

Captures and Wind on 2016-12-01
RhoB+ Male
RhoB+ Mated Females
Wind Speed (m/s) and Direction
Wild Male
Wild Mated Females
Unmated Females
N
NE
E
SE
S
SW
W
NW
50
100
150
200
250
300
350
400
450
0
1
2
3
4
5

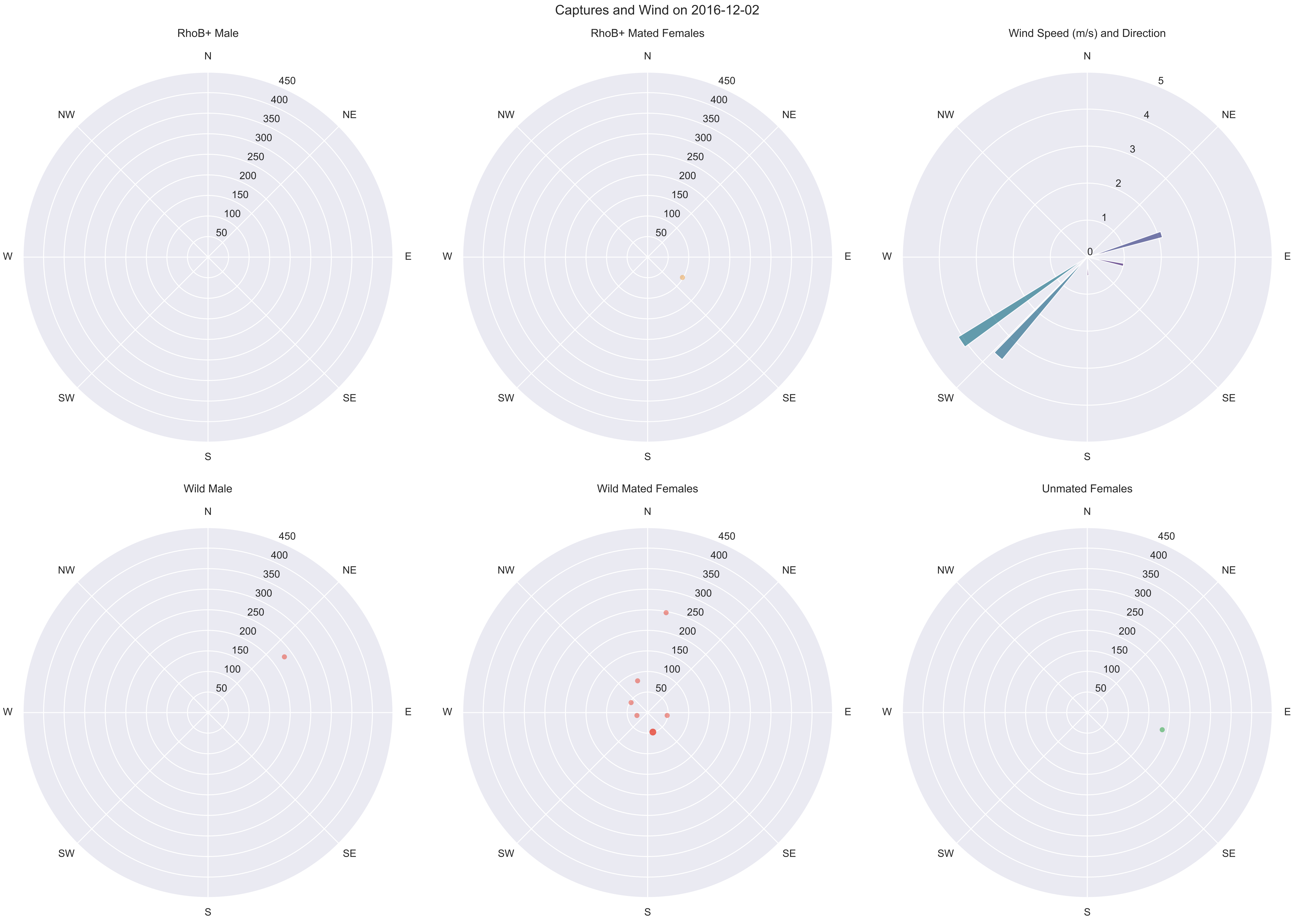
Captures and Wind on 2016-12-02
RhoB+ Male
RhoB+ Mated Females
Wind Speed (m/s) and Direction
Wild Male
Wild Mated Females
Unmated Females
N
NE
E
SE
S
SW
W
NW
50
100
150
200
250
300
350
400
450
0
1
2
3
4
5

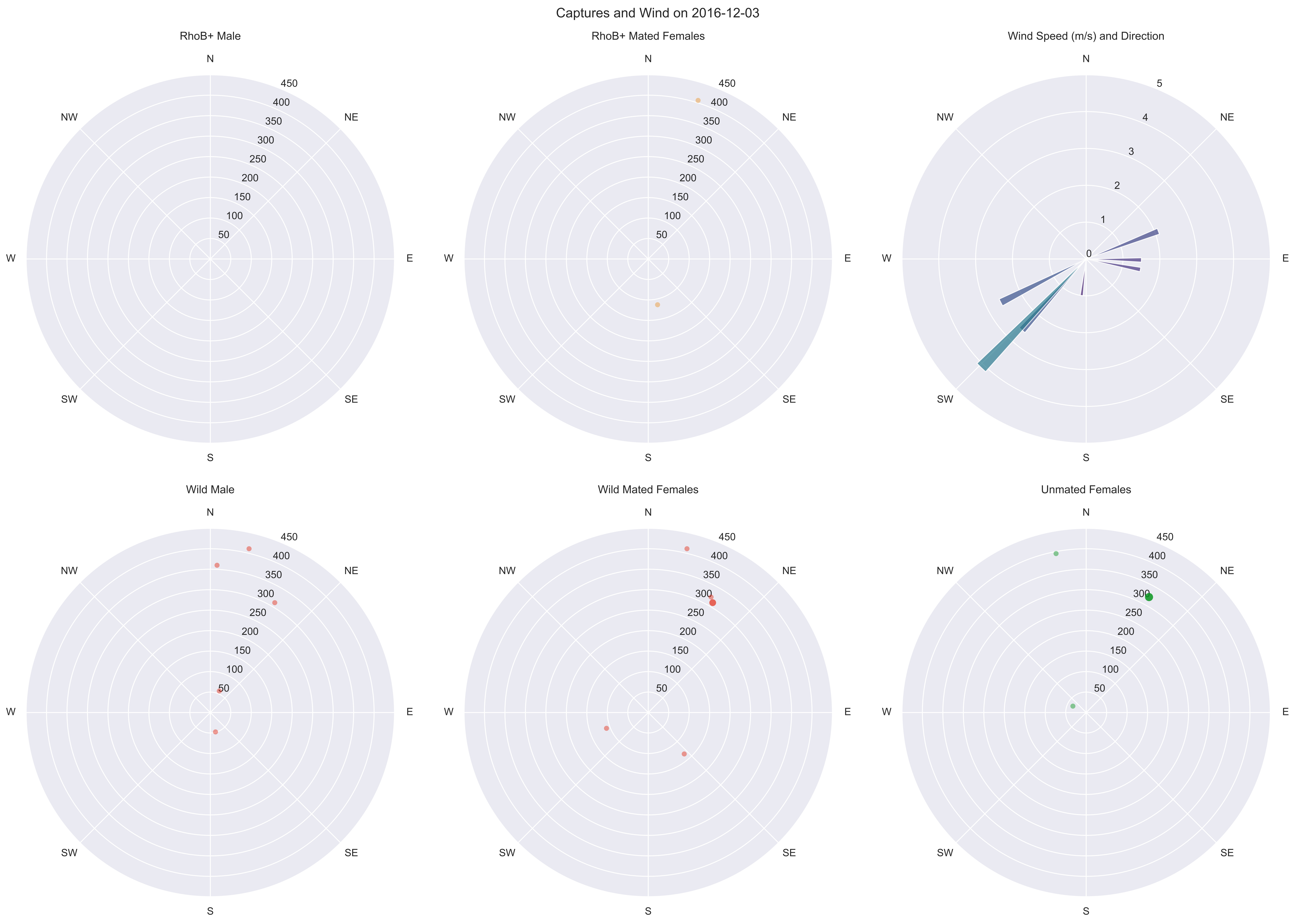

Captures and Wind on 2016-12-03
RhoB+ Male
RhoB+ Mated Females
Wind Speed (m/s) and Direction
Wild Male
Wild Mated Females
Unmated Females
N
NE
E
SE
S
SW
W
NW
50
100
150
200
250
300
350
400
450
0
1
2
3
4
5

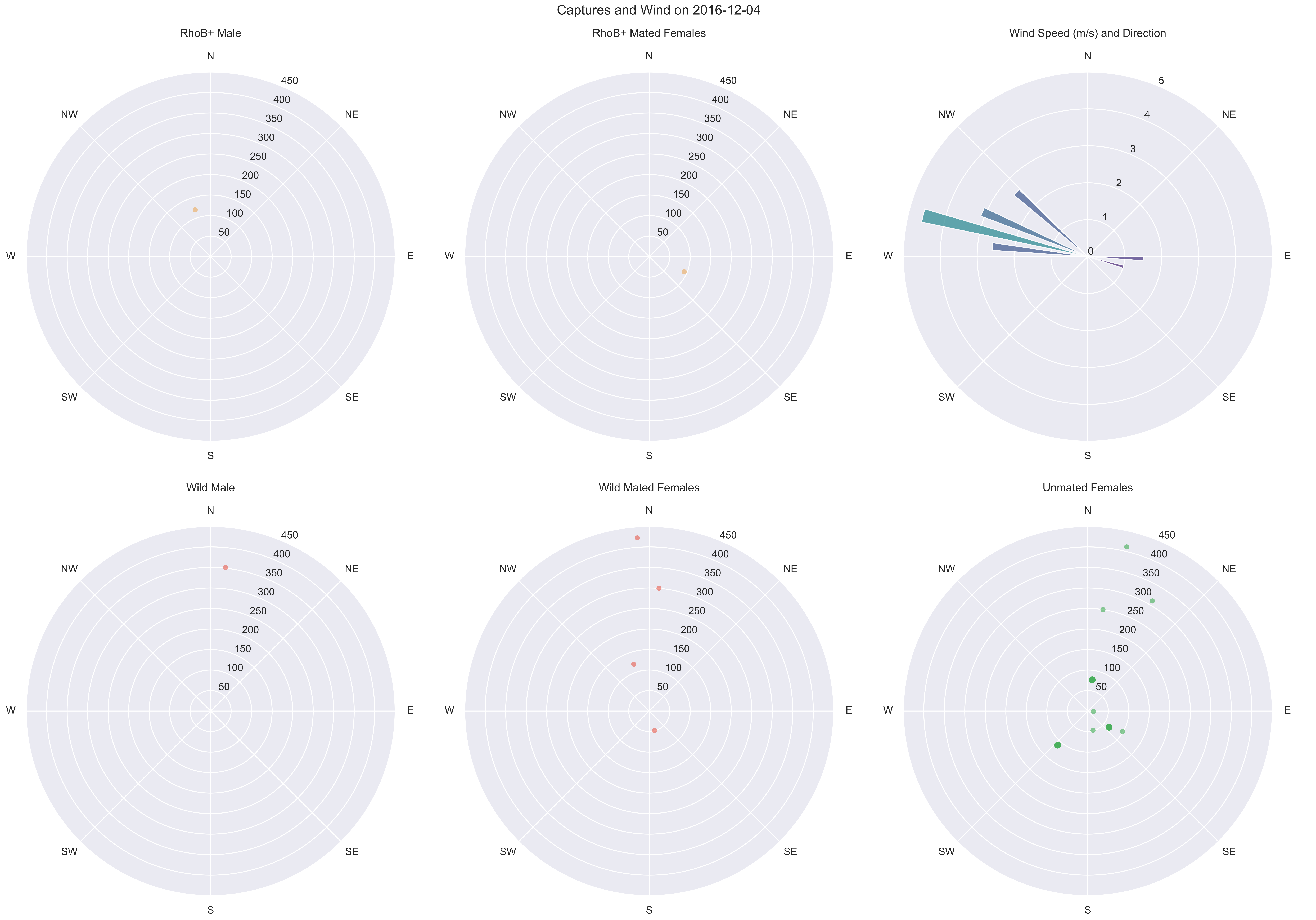
Captures and Wind on 2016-12-04
RhoB+ Male
RhoB+ Mated Females
Wind Speed (m/s) and Direction
Wild Male
Wild Mated Females
Unmated Females
N
NE
E
SE
S
SW
W
NW
50
100
150
200
250
300
350
400
450
0
1
2
3
4
5

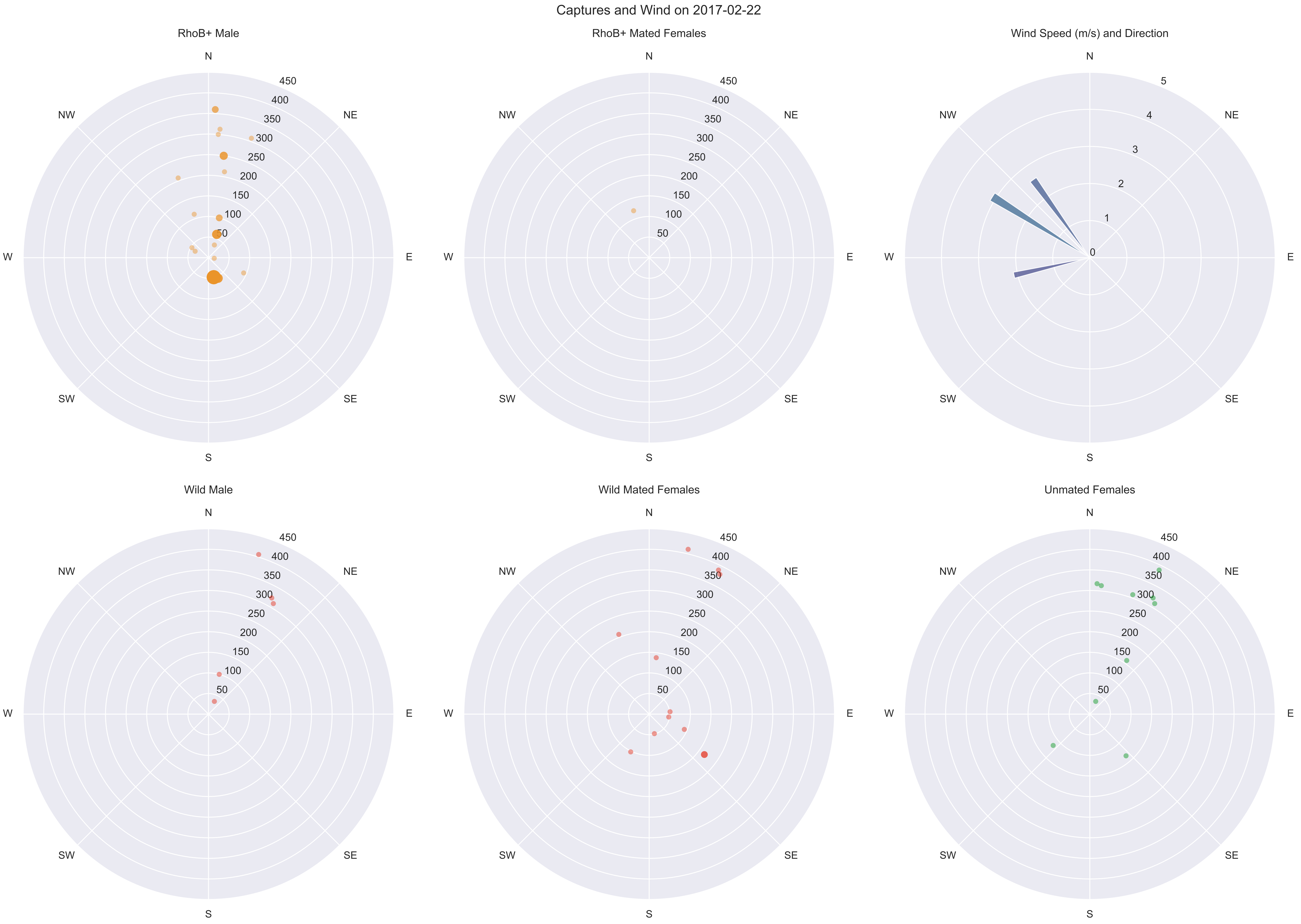
Captures and Wind on 2017-02-22
RhoB+ Male
RhoB+ Mated Females
Wind Speed (m/s) and Direction
Wild Male
Wild Mated Females
Unmated Females
N
NE
E
SE
S
SW
W
NW
50
100
150
200
250
300
350
400
450
0
1
2
3
4
5

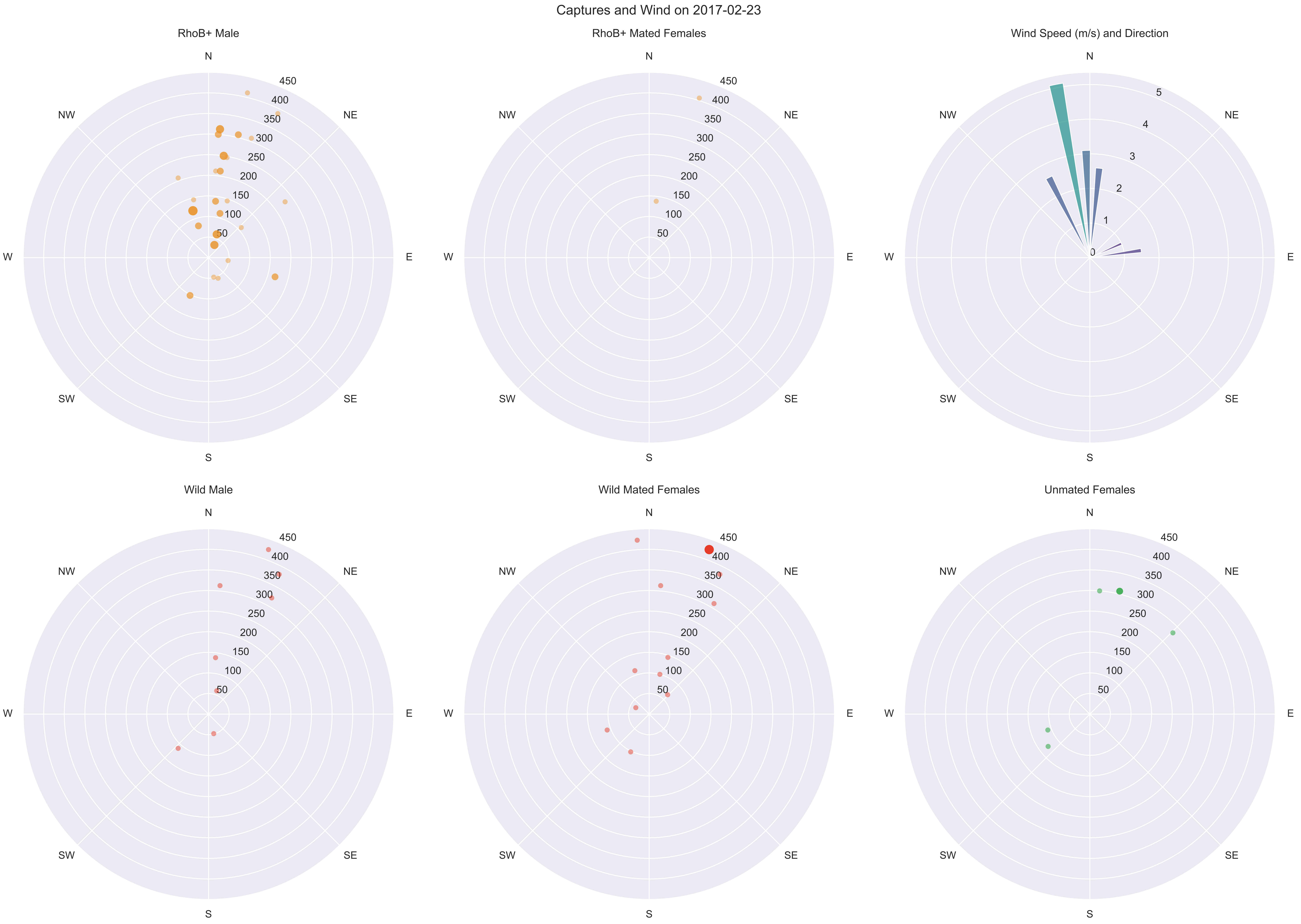
Captures and Wind on 2017-02-23
RhoB+ Male
RhoB+ Mated Females
Wind Speed (m/s) and Direction
Wild Male
Wild Mated Females
Unmated Females
N
NE
E
SE
S
SW
W
NW
50
100
150
200
250
300
350
400
450
0
1
2
3
4
5

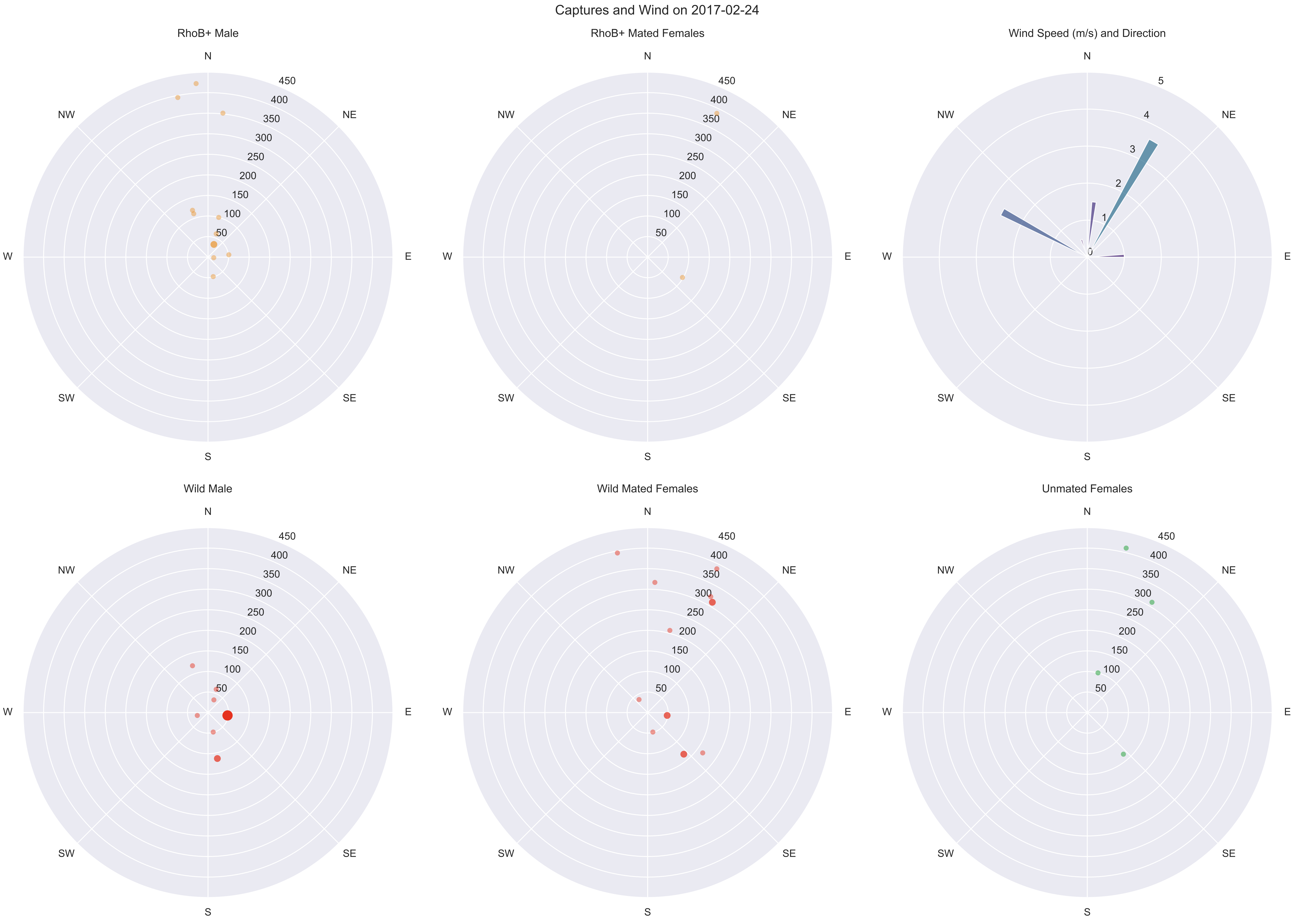
Captures and Wind on 2017-02-24
RhoB+ Male
RhoB+ Mated Females
Wind Speed (m/s) and Direction
Wild Male
Wild Mated Females
Unmated Females
N
NE
E
SE
S
SW
W
NW
50
100
150
200
250
300
350
400
450
0
1
2
3
4
5

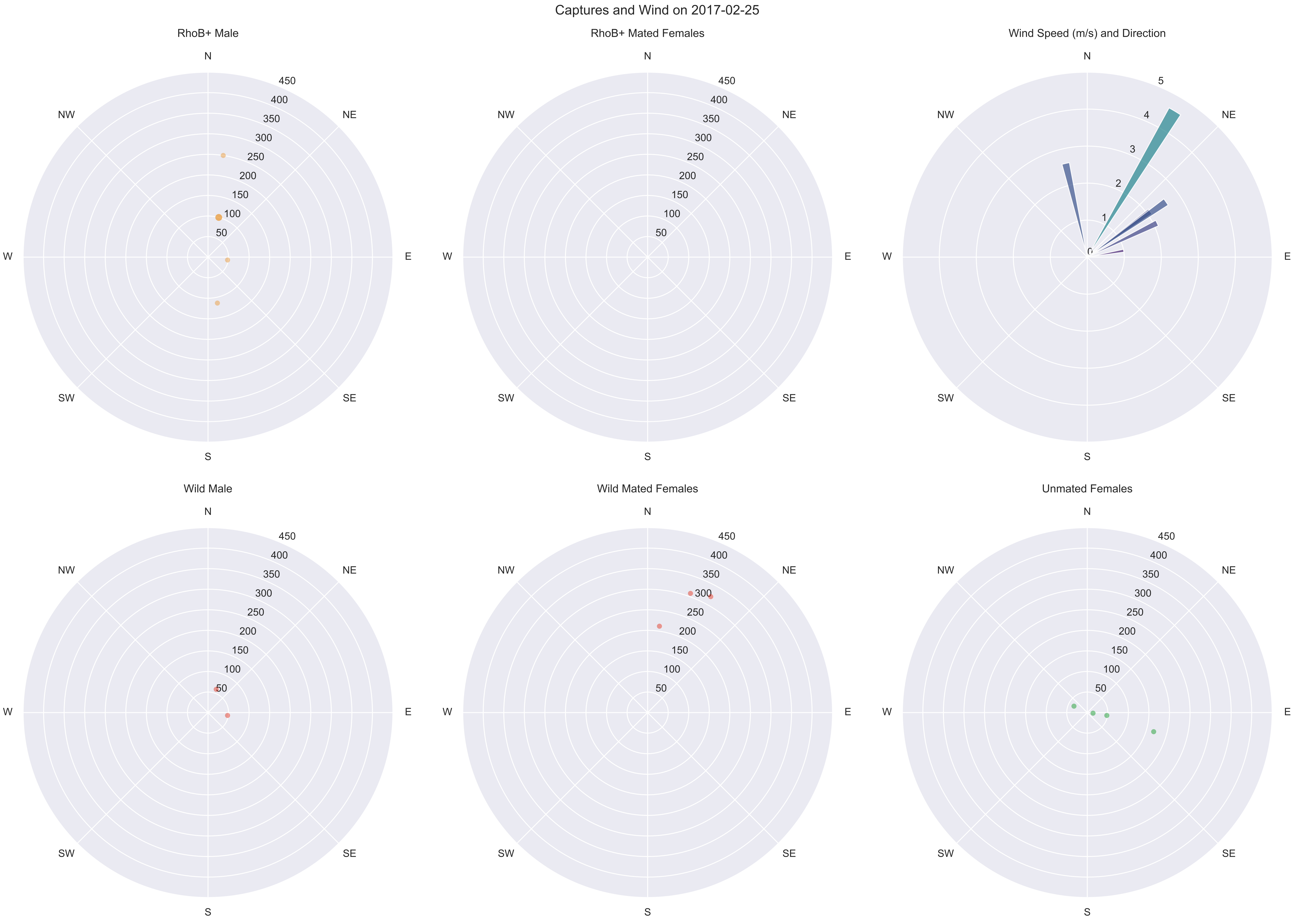

Captures and Wind on 2017-02-25
RhoB+ Male
RhoB+ Mated Females
Wind Speed (m/s) and Direction
Wild Male
Wild Mated Females
Unmated Females
N
NE
E
SE
S
SW
W
NW
450
400
350
300
250
200
150
100
50
5
4
3
2
1
0

Captures and Wind on 2017-02-26

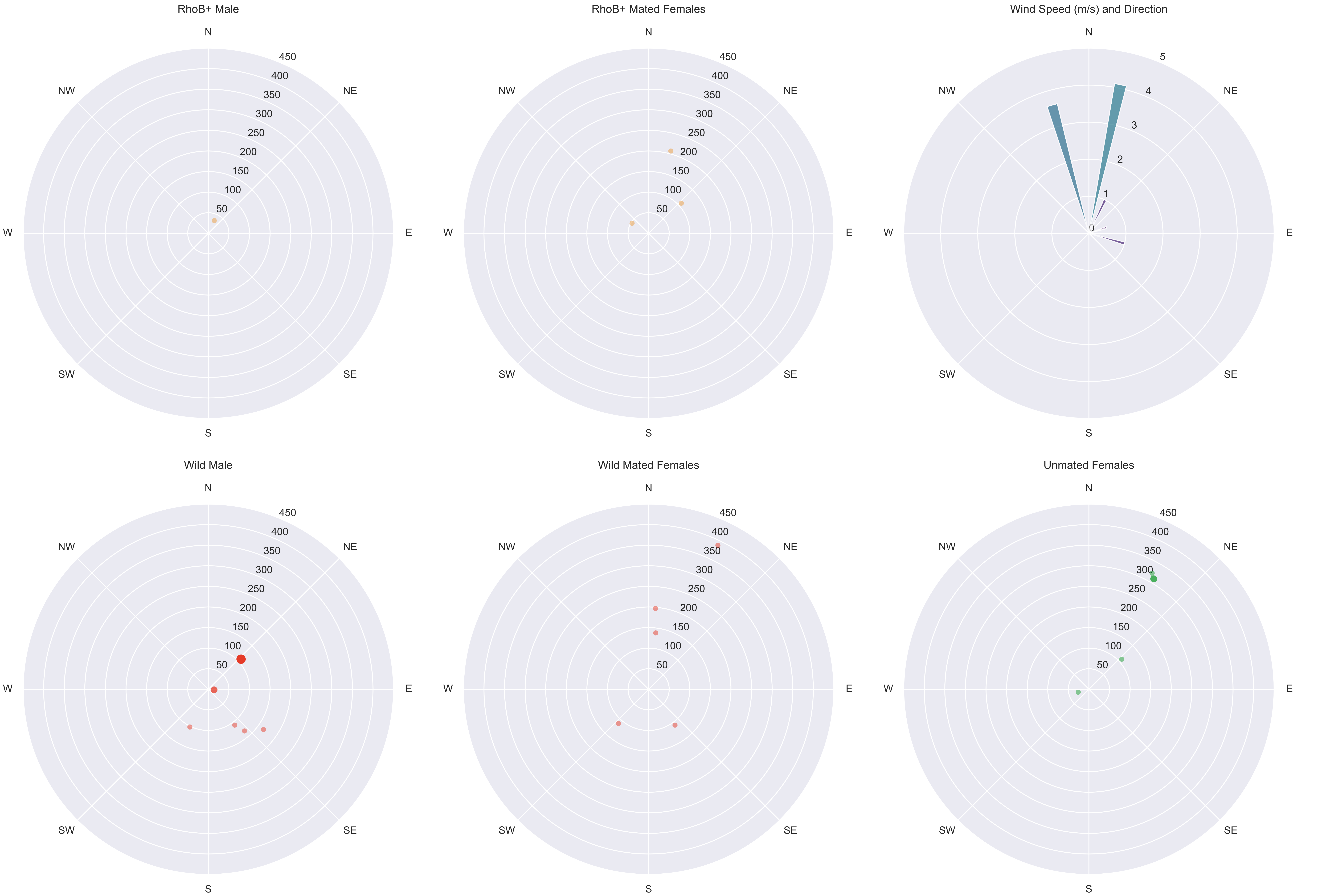

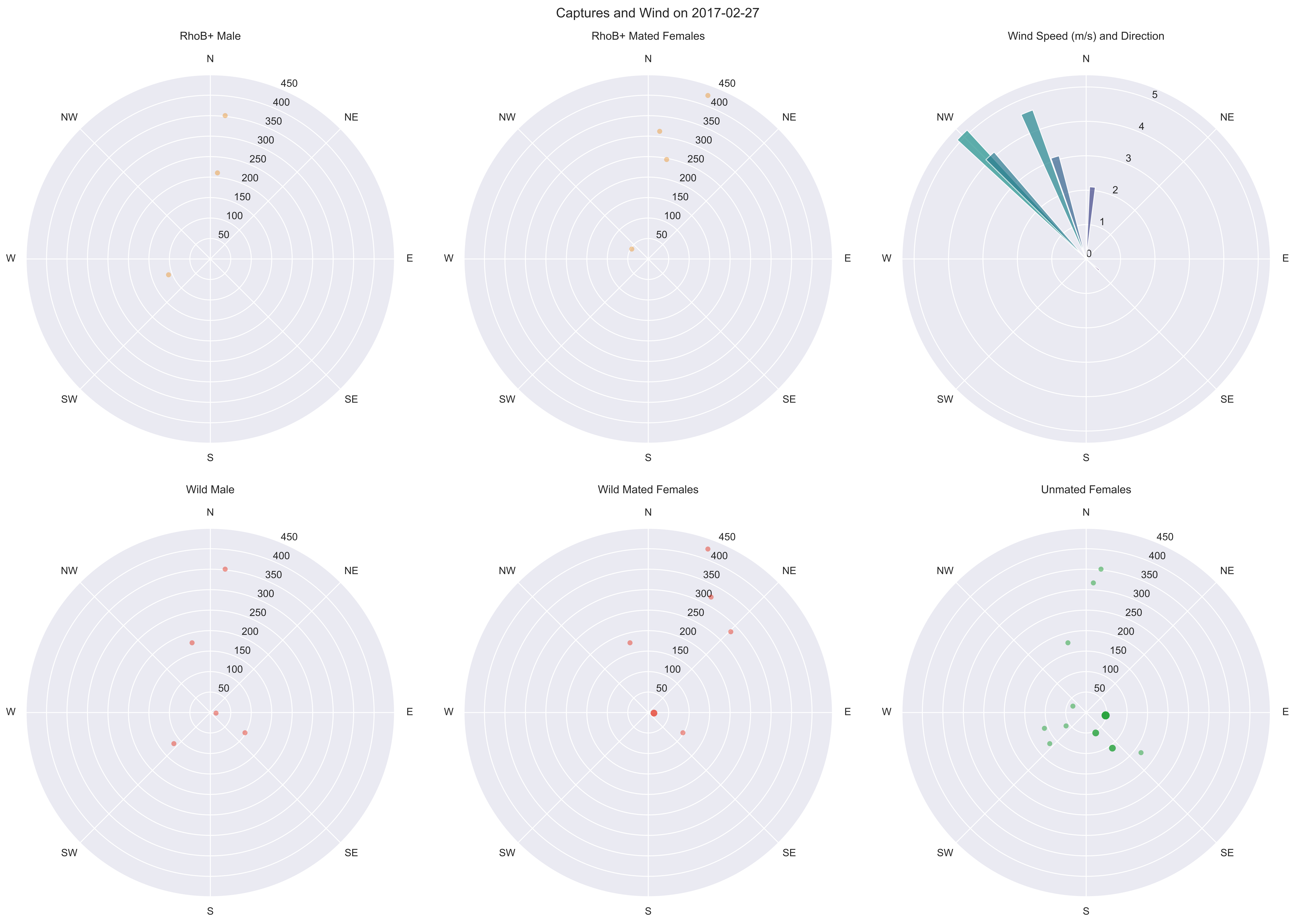

Captures and Wind on 2017-02-27
RhoB+ Male
RhoB+ Mated Females
Wind Speed (m/s) and Direction
Wild Male
Wild Mated Females
Unmated Females
N
NE
E
SE
S
SW
W
NW
50
100
150
200
250
300
350
400
450
0
1
2
3
4
5
